# *DROL1* subunit of U5 snRNP in the spliceosome is specifically required to splice AT–AC-type introns in *Arabidopsis*

**DOI:** 10.1101/2020.10.19.345900

**Authors:** Takamasa Suzuki, Tomomi Shinagawa, Tomoko Niwa, Hibiki Akeda, Satoki Hashimoto, Hideki Tanaka, Yoko Hiroaki, Fumiya Yamasaki, Hiroyuki Mishima, Tsutae Kawai, Tetsuya Higashiyama, Kenzo Nakamura

**Affiliations:** Department of Biological Chemistry, College of Bioscience and Biotechnology, Chubu University, 1200 Matsumoto-cho, Kasugai, Aichi 487-8501, Japan; Cellular and Structural Physiology Institute (CeSPI), Nagoya University, Furo-cho, Chikusa-ku, Nagoya, Aichi 464-8601, Japan; Institute of Transformative Bio-Molecules (WPI-ITbM), Nagoya University, Furo-cho, Chikusa-ku, Nagoya, Aichi 464-8601, Japan; Division of Biological Science, Graduate School of Science, Nagoya University, Nagoya, Aichi 464-8602, Japan; Department of Biological Sciences, Graduate School of Science, The University of Tokyo, 7-3-1 Hongo, Bukyo-ku, Tokyo 113-0033, Japan

**Keywords:** AT–AC-type intron, minor intron, intron splicing, U12-dependent spliceosome, oleosin, abscisic acid response, *Arabidopsis*

## Abstract

An *Arabidopsis* mutant named *defective repression of OLE3::LUC 1* (*drol1*) was originally isolated as a mutant with defects in the repression of *OLEOSIN3* (*OLE3*) after seed germination. In this study, we show that DROL1 is an *Arabidopsis* homolog of yeast DIB1, a subunit of U5 snRNP in the spliceosome. It is also part of a new subfamily that is specific to a certain class of eukaryotes. Comprehensive analysis of the intron splicing using RNA-Seq analysis of the *drol1* mutants revealed that most of the minor introns with AT–AC dinucleotide termini had reduced levels of splicing. Only two nucleotide substitutions from AT–AC to GT–AG enabled AT–AC-type introns to be spliced in *drol1* mutants. Forty-eight genes, including those having important roles in abiotic stress responses and cell proliferation, exhibited reduced splicing of AT–AC-type introns in the *drol1* mutants. Additionally, *drol1* mutant seedlings showed retarded growth, similar to that caused by the activation of abscisic acid signaling, possibly as a result of reduced AT–AC-type intron splicing in the endosomal Na^+^/H^+^ antiporters and plant-specific histone deacetylases. These results indicate that DROL1 is specifically involved in the splicing of minor introns with AT–AC termini, and that this plays an important role in plant growth and development.

**Significance statement:** *Defective Repression of OLE3::LUC 1* (*DROL1)* is a homolog of yeast DIB1, which is a subunit of the U5 snRNP in the spliceosome, but is also part of a new subfamily that is specific to a certain class of eukaryotes. Using RNA-Seq we show that introns with AT–AC dinucleotide termini were specifically retained in the transcriptome of *drol1* mutants and that their splicing plays an important role in plant growth and development.

## Introduction

Dramatic changes in gene expression patterns occur during the developmental transition from seed maturation to seed germination; because the former is characterized by the accumulation of large amounts of storage reserves and the acquisition of dormancy and desiccation tolerance, whereas the latter is characterized by the degradation of storage reserves and the rapid vegetative growth of seedlings. Several seed-specific genes are involved in seed maturation (hereafter referred to as seed-maturation genes), such as the genes encoding seed-storage proteins and oil-body boundary proteins (oleosins), which are turned off or repressed during seed germination and eventually silenced in adult plants. Several important factors that control this dramatic transition in the expression of seed-maturation genes have previously been identified. Several factors for transcriptional regulation and epigenetic modification of gene expression have been identified to control this dramatic transition in the expression of seed-maturation genes, such as PICKLE (Ogas *et al*., 1997), HSI2/VAL1 and HSL1/VAL2 (Suzuki *et al*., 2007; Tsukagoshi *et al*., 2007), ASIL1 (Gao *et al*., 2009), HDA19 (Zhou *et al*., 2013), HDA6, and MEDIATOR (MED) (Chhun *et al*., 2016).

To further identify the factors involved in the repression of seed-maturation genes during germination, we screened *Arabidopsis thaliana* mutants post-germination that were defective in the repression of the *luciferase* (*LUC*) reporter gene expressed under the control of the seed-specific *OLEOSIN3* (*OLE3*) gene promoter (*OLE3::LUC*) (Suzuki *et al*., 2018). An *Arabidopsis* mutant, named as *defective repression of OLE3::LUC 1* (*drol1*), exhibited significantly high and prolonged expression of *OLE3::LUC* after germination when compared with the wild-type (Suzuki *et al*., 2018). The *DROL1* gene encodes a protein homologous to a subunit of the U5 small nuclear ribonucleoprotein particle (snRNP) that is involved in pre-mRNA splicing. However, the precise role of DROL1 in splicing and how the defect in a splicing factor causes the derepression of *OLE3* during germination remains unknown.

Many eukaryotic genes contain introns, which are excised from pre-mRNAs via a complex process mediated by the spliceosome to form functional mRNAs. The terminal GT–AG dinucleotides are the most conserved intron structures in all eukaryotes, and these canonical GT–AG introns are spliced by the major U2-dependent spliceosome, containing U1, U2, U4, U5, and U6 snRNAs and approximately 80 proteins (Wilkinson *et al*., 2019). In addition to the U2-dependent splicing of the canonical GT–AG introns, a small number of unusual introns containing non-consensus AT–AC termini and highly conserved sequences at the 5′ splice site and the branch point are spliced by the minor U12-dependent spliceosome (Jackson, 1991; Tarn and Steitz, 1996a; Tarn and Steitz, 1996b; Turunen *et al*., 2013). The U12-dependent spliceosome consists of four unique snRNAs, U11, U12, U4_atac_, and U6_atac_, and only the U5 snRNA is shared with the U2-dependent spliceosome (Tarn and Steitz, 1996a; Tarn and Steitz, 1996b). The protein composition of each snRNP of the U12-dependent spliceosome is very similar, if not identical, to that of the U2-dependent spliceosome (Schneider *et al*., 2002; Will *et al*., 1999).

While U12 spliceosome-dependent splicing was first identified for AT–AC introns, it was later found that GT–AG is the more common terminal dinucleotide even for U12-type introns. The U12-type introns have been identified in various eukaryotes and represent ≤0.2% of all pre-mRNA introns (Burge *et al*., 1998; Szcześniak *et al*., 2013; Zhu and Brendel, 2003). However, unlike U2-type introns, U12-type introns are absent in certain species, such as nematodes (*Caenorhabditis elegans*) and budding yeast (*Saccharomyces cerevisiae*). The U12-type introns seem to be enriched in genes within certain functional classes including DNA replication and repair, transcription, RNA processing, and translation, and these genes usually contain only one U12-type intron (Turunen *et al*., 2013; Burge *et al*., 1998). The splicing of U12-type introns appears to be significantly slower than that of U2-type introns, and it has been suggested that U12-type introns play a role in regulating the expression of specific sets of genes (Turunen *et al*., 2013).

In this study, we have shown that DROL1 is a homolog of yeast DIB1, a protein subunit of U5 snRNP, and that the mutation of *DROL1* causes the specific retention of AT–AC-type introns but not GT–AG-type introns. This increased retention of AT–AC-type introns upregulates the expression of abscisic acid (ABA)-responsive genes and delays germination and seedling growth. These results indicate that the splicing of minor AT–AC-type introns plays an important role in plant growth and development.

## Results

### DROL1 is homologous to yeast DIB1 but is also comprised of a new subfamily

All eukaryotic genomic databases searched in this study contained *DROL1* homologs, but we found no homologous genes in the prokaryotic genomic databases. We selected *DROL1* gene homologs of *A. thaliana*, rice (*Oryza sativa*), bryophyte (*Physcomitrella patens*), green alga (*Chlamydomonas reinhardtii*), human (*Homo sapiens*), mouse (*Mus musculus*), nematode (*Caenorhabditis elegans*), fission yeast (*Schizosaccharomyces pombe*), and budding yeast (*Saccharomyces cerevisiae*) to use in this investigation. The predicted amino acid sequences of their DROL1 homologs were aligned (Figure 1a) and used for phylogenetic analysis (Figure 1b).

**Figure 1.**
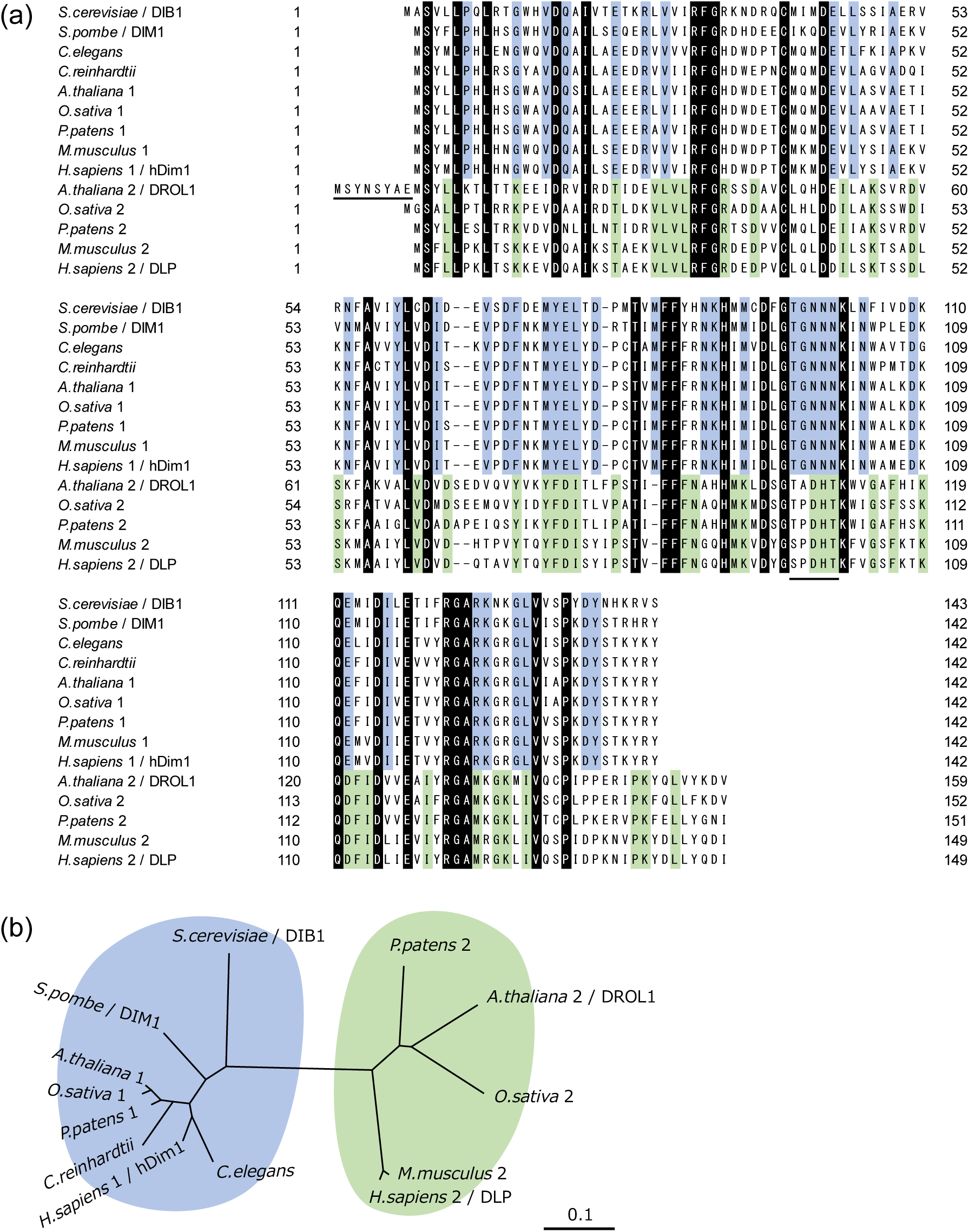
Comparison of DROL1 homologs. (a) Alignment of amino acid sequences of fourteen DROL1 homologs from nine model organisms. The species name is shown only if it harbors DIB1 subfamily members, and the names were followed by the numbers 1 and 2 to indicate if the DIB1 and DROL1 subfamily members were present, respectively. Gene IDs and source organisms were as follows: AT5G08290.1, AT3G24730.1 (DROL1): *Arabidopsis thaliana*; Os07t0202300-01, Os11t0297900-01: *Oryza sativa*; Pp1s141_103V6.1, Pp1s12_196V6.1: *Physcomitrella patens*; Cre01.g055416.t1.1: *Chlamydomonas reinhardtii*; TXN4A_MOUSE, TXN4B_MOUSE: *Mus musculus*; TXN4A_HUMAN, TXN4B_HUMAN: *Homo sapiens*; Y54G2A.75.1: *Caenorhabditis elegans*; SPCC16A11.05c: *Schizosaccharomyces pombe*; and YPR082C: *Saccharomyces cerevisiae*. The conserved amino acid residues among all homologs and the DIB1 and DROL1 subfamilies are indicated in black, blue, and green, respectively. (b) Phylogenetic trees of the DROL1 homologs. Mouse (TXN4A_MOUSE) and human (TXN4A_HUMAN) DROL1 orthologs have the same amino acid sequences, and consequently, the node corresponding to mouse DROL1 was omitted.

The multiple sequence alignment revealed that the DROL1 homologs could be divided into two subfamilies. One subfamily was represented by the *S. cerevisiae* DIB1, a subunit of the U5 snRNP (Gottschalk, 1999; Stevens and Abelson, 1999), and this is hereafter referred to as the DIB1 subfamily. All members of the DIB1 subfamily contained 142 amino acids (aa), except DIB1 (143 aa), and shared >80% sequence identity. All eukaryotes examined in this study contained at least one DIB1 homolog. The other subfamily was comprised of DROL1 and its close homologs and is hereafter referred to as the DROL1 subfamily. The DROL1 subfamily proteins were variable in length, ranging from 149 to 159 aa, and shared less sequence identity among each other than the DIB1 subfamily members; for example, *A. thaliana* DROL1 and *H. sapiens* DLP share approximately 60% sequence identity. Unlike members of the DIB1 subfamily, certain eukaryotes, such as *S. pombe*, *S. cerevisiae*, *C. elegans*, and *C. reinhardtii*, do not contain any proteins belonging to the DROL1 subfamily. Further analysis showed that genomes of other eukaryotes, including frog (*Xenopus laevis*), slugfish (*Branchiostoma floridae*), sea urchin (*Strongylocentrotus purpuratus*), octopus (*Octopus bimaculoides*), plasmodial slime mold (*Physarum polycephalum*), and oomycetes (*Globisporangium ultimum* and *Phytophthora kernoviae*), encode both DIB1 and DROL1 subfamily members, whereas the fruit fly (*Drosophila melanogaster*), western honey bee (*Apis mellifera*), sea squirt (*Ciona intestinalis*), and cellular slime mold (*Dictyostelium discoideum*) encode only DIB1 subfamily members (Figure S1). This distribution of the DROL1 subfamily in organisms is almost consistent with that of the U12-dependent spliceosomes, except insects and sea squirt, which are reported to harbor U12-dependent spliceosomes but not DROL1 orthologs (Bai *et al*., 2019; Bartschat and Samuelsson, 2010; Turunen *et al*., 2013).

### Translational initiation site of *DROL1*

The multiple sequence alignment of the DROL1 subfamily members indicated that DROL1 has a longer N-terminal amino acid sequence than other subfamily members (underlined in Figure 1a). It also contained a second ATG codon 24 bp downstream of the first ATG, and translational initiation from the second ATG produced a protein like that of the other subfamily members. To confirm the 5′ end of the *DROL1* mRNA, we performed 5’ rapid amplification of the cDNA ends (5′ RACE). The cDNA fragments that were amplified by 5′ RACE were ligated to adapters and sequenced using next-generation sequencing (NextSeq500, Illumina). The sequence reads obtained were then aligned to the *DROL1* cDNA sequence, and four major 5′ ends (T1–T4) were identified (Figure S2a). The most upstream transcription start site (T1) was located only 6 bps upstream of the first ATG (Figure S2a, orange arrowhead). The second major site (T2) was located between the two ATG codons and 16 bps upstream of the second ATG codon (Figure S2a, red arrowhead). The remaining two transcription start sites (T3 and T4) were located downstream of the second ATG, implying that these transcripts represented degradation products.

To test which ATG codon is important for the functional expression of *DROL1*, we constructed four mutant variants of the *DROL1* gene (1AATG, 1ATGA, 2AATG, and 2ATGA), wherein one adenine (A) nucleotide was inserted upstream or downstream of the first and second ATG codons, respectively. When the A nucleotide was cloned upstream of the translation initiation site of *DROL1*, the gene was predicted to maintain its function and complement the *drol1-1* mutant phenotype. Three mutant variants of *DROL1* (1AATG, 1ATGA, and 2AATG) with insertions of A upstream of the second ATG complemented the *drol1-1* mutant phenotype (Figure S2b). In contrast, the gene with the A insertion immediately after the second ATG, namely 2ATGA, failed to complement the *drol1-1* mutant phenotype (Figure S2b). These results and those of the multiple sequence alignment suggest that the translation initiation of *DROL1* occurs from the second ATG and not the first.

### Introns requiring DROL1 for splicing

Since *Arabidopsis* DROL1 is a homolog of *S. cerevisiae* DIB1 which is a subunit of the U5 snRNP, defects in *DROL1* are predicted to directly affect intron splicing. To identify the changes in the intron splicing patterns of the *drol1* mutant, we attempted to estimate the intron retention levels in the mRNAs with RNA-Seq. The original *drol1-1* mutant carries a point mutation at the splice site of the third intron of *DROL1* gene that interferes with its correct splicing, while the *drol1-2* mutant carries a T-DNA insertion in the second intron of *DROL1* gene (Suzuki *et al*., 2018). Total RNA was extracted from the wild-type, *drol1-1*, and *drol1-2* seedlings 3 days after sowing and was used to construct cDNA libraries for next-generation sequencing in three independent experiments. Single-end libraries were constructed using 86-bps sequences, and approximately 70 million reads (on average) were obtained per library (Table S1). The short reads were mapped onto the *Arabidopsis* genome sequence using Tophat (Trapnell *et al*., 2009) and intron splicing patterns were analyzed using ASpli (Estefania *et al*., 2021).

Of the approximately 125,000 introns in the *Arabidopsis* genome, 2,741 were further analyzed as there were enough supporting reads to compare them between the wild-type and *drol1-1*. The percent of intron retention (PIR) was calculated and compared (Figure 2a and S3). The PIR was similar for most of the Col and *drol1-*1, and the correlation coefficient was high (0.94), which suggested that most introns in *drol1-1* were spliced in a similar manner in the wild-type. Of the 2,741 introns, 182 showed significantly different splicing patterns (false discovery rage [FDR] < 0.01; large blue and red dots in Figure 2a). We searched for the shared features among the 182 introns and found that one fourth (49/182; red dots in Figure 2a) had nucleotide sequences that started with AT and ended with AC, hereafter referred to as AT–AC-type introns. This proportion of AT–AC-type introns (27%) was far greater than the proportion in the *Arabidopsis* genome (0.06%; *p* < 10^−30^ with chi-square test).

**Figure 2.**
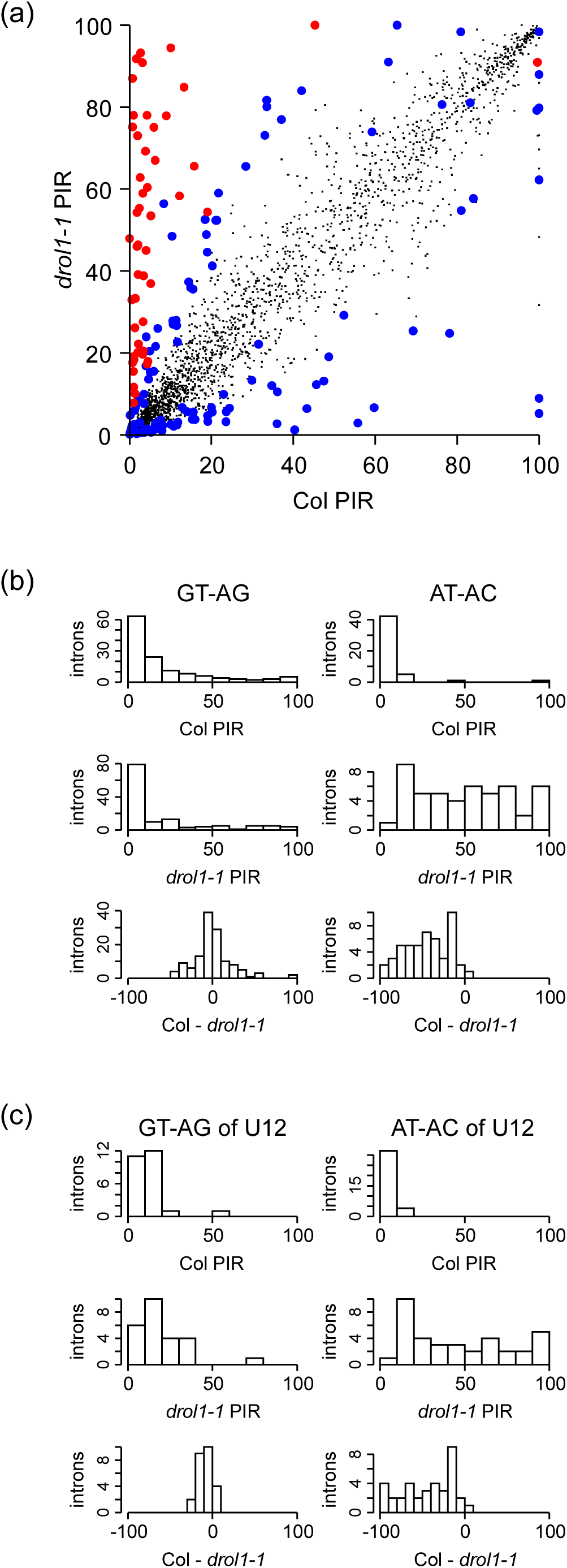
Global analysis of intron retention in the *drol1-1* mutant. (a) Scatter plot of the percent intron retention (PIR) for the wild-type (Col) and *drol1-1* mutant. All small and large dots represent 2,741 introns which were supported by enough short reads for the analysis by ASpli (Estefania *et al*., 2021). AT–AC-type introns and other types of intron (mostly GT–AG) with FDR < 0.01 are plotted as red and blue large dots, respectively. (b) Histogram of PIR. Counts of GT–AG- and AT–AC-type introns grouped by their PIR are plotted in the left and right columns, respectively. The first and second rows represent the observed PIR in the wild-type and *drol1-1*, respectively. The third row represents the differences in PIR (*drol1-1* was subtracted from Col) between the wild-type and *drol1-1*. (c) Histograms of PIR for 25 GT–AG- and 33 AT–AC-type of U12-dependent introns. Similar experiments were performed for *drol1-2* (Figure S4).

Most of the GT–AG- and AT–AC-type introns in the 182 differentially spliced introns were almost completely spliced in the wild-type (Figure 2b, top row). Similar patterns were observed for the GT–AG-type introns in the *drol1-1* mutant (Figure 2b, left middle panel). The difference in the PIR between the wild-type and *drol1-1* mutant (Figure 2b, bottom left panel) was concentrated near the center of the histogram, indicating that most PIR of the GT–AG-type introns remained unaffected by the DROL1 deficiency. However, the PIR of the AT–AC-type introns varied in the *drol1-1* mutant (Figure 2b, right middle panel), and almost all differences were negative (Figure 2b, right bottom panel), indicating that most PIRs of the AT–AC-type introns was increased in the *drol1-1* mutant. These results suggest that DROL1 is specifically required for the splicing of AT–AC-type introns, although the dependence on DROL1 for splicing varied among introns. Similar tendencies were also observed in the 3-day-old seedlings of the *drol1-2* mutants (Figure S4).

All AT–AC-type introns that were significantly retained in the *drol1* mutants are listed in Table 1, and this includes 49 introns belonging to 48 genes, which represent candidates that may require DROL1 for their splicing.

### Intron retention in *histone deacetylase 2B* (*HD2B*) mRNA in *drol1* mutants

Among these 182 introns, the third intron of the *HD2B* gene, a member of the *HD2* gene family encoding plant-specific histone deacetylases (Hollender and Liu, 2008), was most significantly retained in 3-day-old *drol1-1* mutant plants (Table 1).

The compiled reads mapped into the *HD2B* gene in the wild-type and *drol1* mutant seedlings are presented in Figure 3(a)–(c). Short reads obtained from 3-day-old wild-type seedlings were only mapped into the exons (Figure 3a and d). However, a significant number of short reads were mapped to the third intron of the *HD2B* gene in *drol1-1* and *drol1-2* mutants, which was an AT–AC-type (Figure 3b–d). By contrast, the other seven introns of the *HD2B* with canonical terminal sequences were completely spliced. These results indicate the retention of AT–AC-type introns in a substantial number of *HD2B* transcripts in *drol1* mutants.

**Figure 3.**
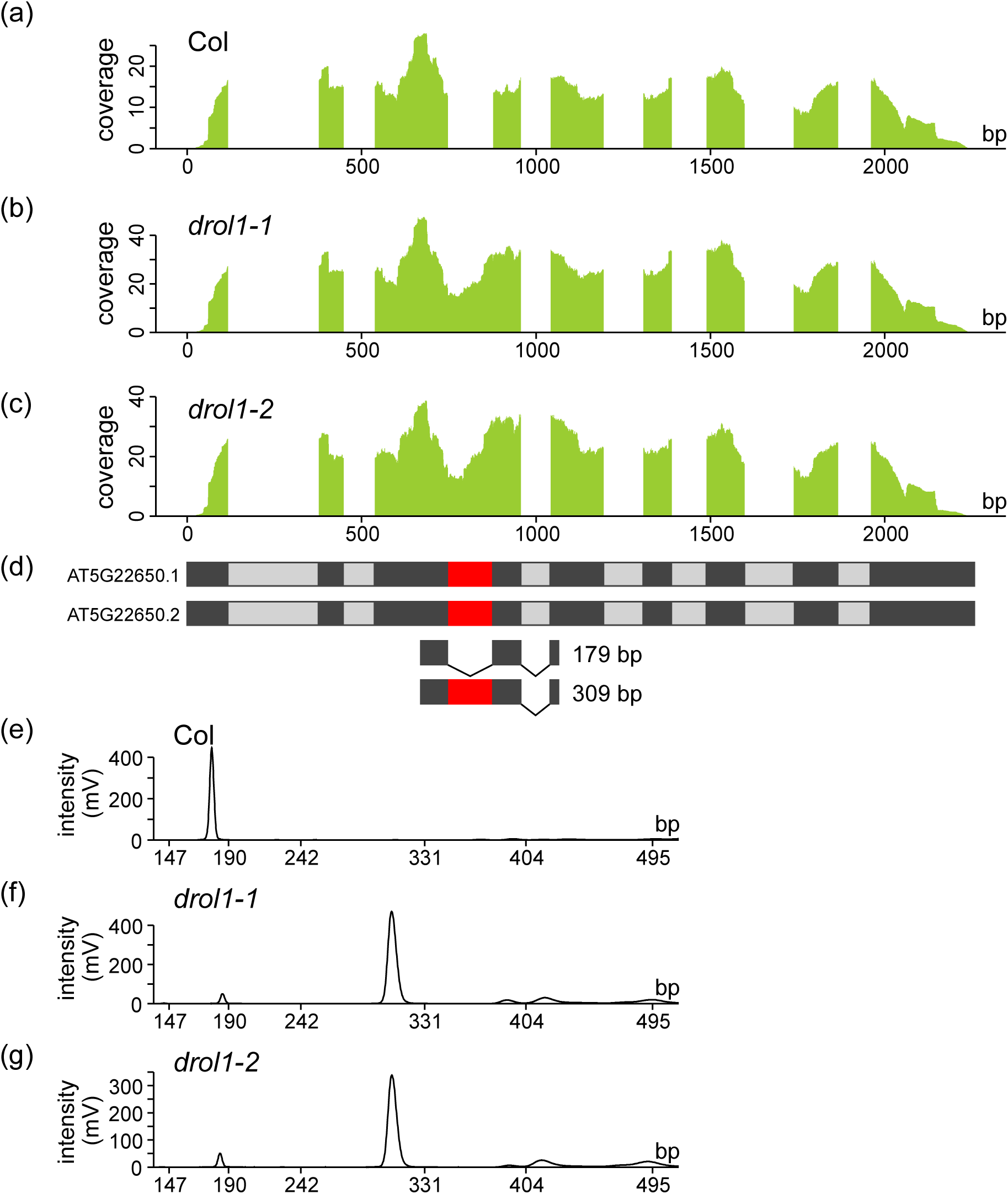
RNA-Seq and RT-PCR analysis for the retention of the third intron of *HD2B* in the wild-type and *drol1* mutants. (a–c) RNA-Seq analysis of *DROL1*. Total RNAs extracted from 3-day-old wild-type (Col) (a), *drol1-1* (b), and *drol1-2* (c) seedlings that were subjected to cDNA library construction for next-generation sequencing. The reads obtained were aligned to the *HD2B* gene. The x-axis represents the reference sequences, and the y-axis indicates the coverage of the short reads. (d) Gene structure of *HD2B*. Black and gray boxes represent exons and introns, respectively. Red boxes indicate AT–AC-type introns. The only difference between AT5G22650.2 and AT5G22650.1 is a 3 bps truncation of the 5′ termini in the eighth exon. The expected RT-PCR products in (e–g) were also indicated with their sizes. (e–g) Chip electrophoresis for the DNA fragment amplified by RT-PCR from the wild-type (e), *drol1-1* (f), and *drol1-2* (g) RNA. The x-axis represents the size of the DNA in bp, and the y-axis indicates the intensity of fluorescence due to the DNA. Similar experiments were performed for *HD2C*, *HD2D*, *NRPA2*, and *NHX5* (Figure S5–S8).

The retention of the AT–AC-type intron in the *HD2B* gene in *drol1* mutants was also confirmed by RT-PCR. From the wild-type RNA, a single cDNA fragment of 179 bp corresponding to the completely spliced mRNA was amplified with primers designed for the third and fifth exons (Figure 3d and e). However, from the *drol1* mutants, the cDNA fragments of 309 bp for the mRNA that had third introns with AT–AC termini were recovered (Figure 3f and g). Similar results were obtained for the AT–AC-type introns of *HD2C*, *HD2D*, *NRPA2,* and *NHX5* genes (Figure S5–S8). These results further indicated that DROL1 is specifically required for the splicing of AT–AC-type introns.

### DROL1 is required for the splicing of only AT–AC-type introns but not other types of the U12-dependent introns

Non-canonical AT–AC-type introns are spliced by the minor U12-dependent spliceosome. However, most of the U12-dependent introns contain GT–AG dinucleotide termini, with consensus sequences that are different from those of the canonical U2-dependent introns (Turunen *et al*., 2013). Therefore, we analyzed the function of DROL1 in the splicing of U12-dependent introns. According to the U12 Intron Database (U12DB; http://genome.crg.es/datasets/u12/), the *Arabidopsis* genome contains 246 U12-type introns, of which 58 (25 for GT–AG-type and 33 for AT–AC) were present in our analyzed data. Most of the U12-dependent introns had PIRs of nearly zero in the wild-type, independent of the terminal dinucleotides (Figure 2c, top row). Although the GT–AG-type U12-dependent introns showed similar PIRs in *drol1-1* and *drol1-2*, many AT–AC-type U12-dependent introns exhibited increased PIRs in *drol1-1* and *drol1-2* (Figure 2c and S4c, middle row). The differences in the PIRs between the wild-type and *drol1* mutants also indicated a specific increase of the PIR with the AT–AC-type introns in *drol1-1* and *drol1-2* (Figure 2c and S4c, bottom row). These results are consistent with the analysis of the differentially retained introns (Figure 2a and b) and indicated that DROL1 was required only for the splicing of AT–AC-type introns, not U12-dependent introns.

### Reduced retention rate of the AT–AC-type introns in *drol1* after conversion to GT–AG-type

The third intron of *HD2B* is a U12-dependent AT–AC-type intron, and the PIR of this intron was higher in *drol1-1* and *drol1-2* mutants than in the wild-type (dPIR = 86.3 in Table 1). The entire sequence of the *HD2B* gene, including the promoter was cloned, and the AT–AC terminal dinucleotide of the third intron was replaced by GT–AG (Figure 4a). The resulting mutant variant of *HD2B* was introduced into the *drol1-2* mutant, and two independent transformants, HD2BgmI/*drol1-2* #1 and #2, were obtained. The total RNA was extracted from 5-day-old seedlings of both transformants, and the PIR values were measured using RNA-Seq. Although the PIRs of the endogenous AT–AC-type introns (two right gray bars) were almost the same as the non-transformed *drol1* mutants, those of the GT–AG-type introns transcribed from the transgene (two right white bars) were significantly decreased (*p* < 0.05 with Student’s t-test; Figure 4b). Similar results were also obtained for the AT–AC-type intron of *HD2C*, a homolog of *HD2B* (Figure 4c). These results indicate that the mutation of AT–AC to GT–AG in the U12-dependent introns abolished their requirement for DROL1 for splicing and clarified its specific role in the splicing of AT–AC-type introns.

**Figure 4.**
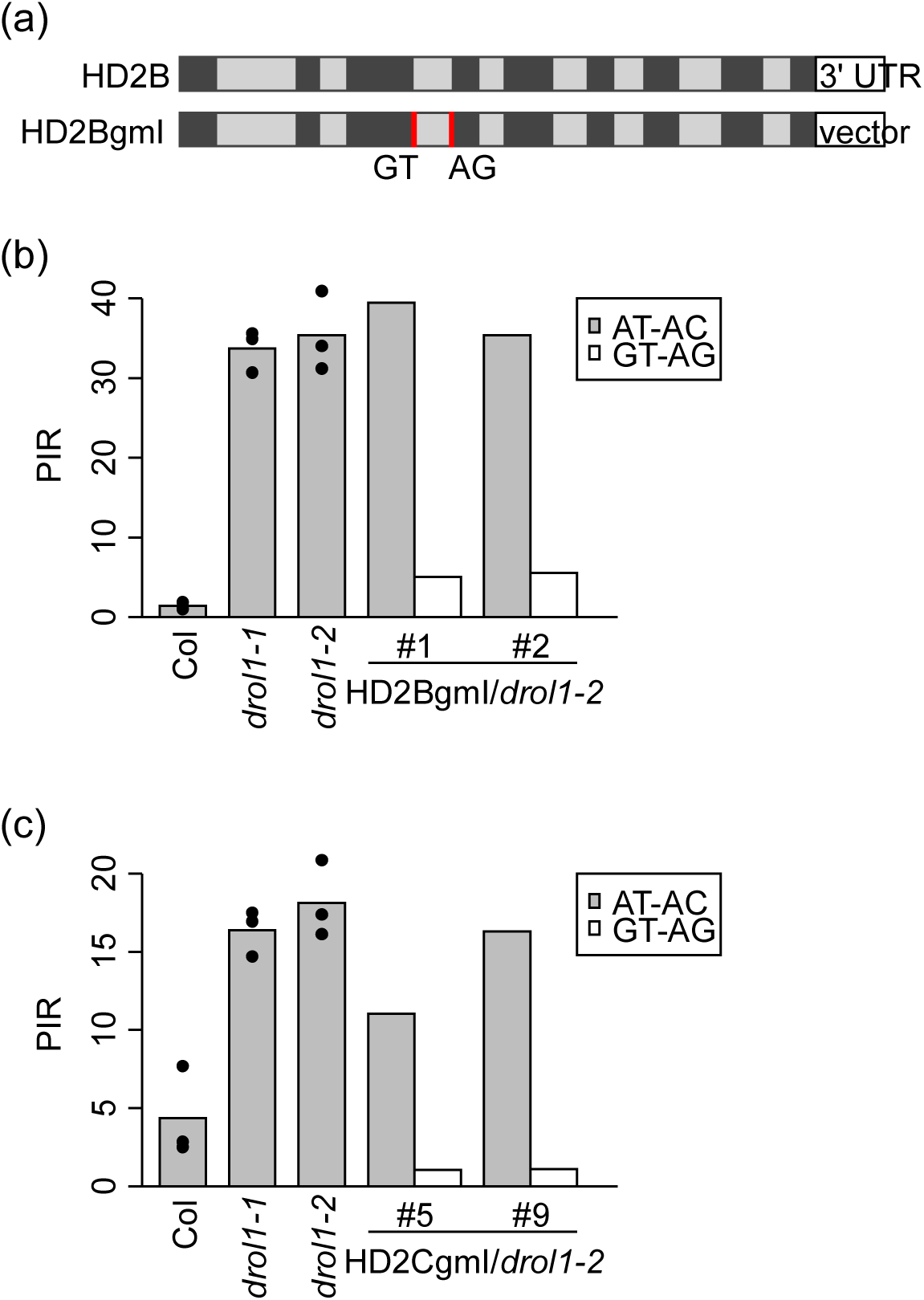
Retention rates of introns converted from the AT–AC-type to the GT–AG-type in *drol1-2*. (a) Gene structure of the endogenous and modified *HD2B*. Nine exons and the 3′ untranslated region (3′ UTR) in exon 9 are shown as black and white boxes, respectively. Two-point mutations, which were used to convert AT–AC-type introns to GT–AG-type introns, are indicated by red bars. *HD2BgmI* is recombined to vector at the stop codon. (b) Percent intron retention (PIR) of the third intron of *HD2B*. The PIRs of the AT–AC-type and GT–AG-type third intron in strains are shown as gray and white bars, respectively. Reads overlapping the third junction of exons from HD2BgmI/*drol1-2* were proportionally distributed to endogenous *HD2B* and transformed *HD2BgmI* based on reads for 3′ UTR and vector sequences followed by a stop codon, respectively. (c) PIR of the third intron of *HD2C*. The experimental scheme was same as for *HD2B*.

### Comprehensive transcriptomic analysis of *drol1* mutants

Genome-wide expression patterns were compared between the wild-type and *drol1* mutants. Total RNA was additionally extracted from 5- and 7-day-old seedlings of the wild-type, *drol1-1*, and *drol1-2*. All short reads, including those from the 3-day-old seedlings, were aligned with the *Arabidopsis* cDNA and counted by transcript. The data were analyzed using edgeR (Robinson *et al*., 2010), and differentially expressed genes (DEGs) were selected (FDR < 0.001). Compared with the wild-type, 1,344 and 2,762 genes were upregulated and downregulated in the 3-day-old *drol1-1* mutant seedlings, respectively (Figure S9). There were fewer DEGs in the *drol1-2* mutant than in the *drol1-1* mutant, although most corresponded with those in the *drol1-1* mutants (88%–90%), suggesting that the effects of *drol1-2* were weaker than those of *drol1-1* on the transcriptome. Similar patterns were observed in the 5- and 7-day-old seedlings, although the DEGs were decreasing (Figure S9).

To characterize the upregulated and downregulated genes in the *drol1* mutants, we performed Gene Ontology (GO) analysis using PANTHER (http://go.pantherdb.org/). Of the 374 genes commonly upregulated in the *drol1-1* and *drol1-2* mutants (Table S2), those related to “lipid storage,” “response to abscisic acid,” “response to water deprivation,” and “response to salt stress” were overrepresented (Table S3). In the 3-day-old *drol1* seedlings, the upregulation of genes involved in the response to dehydration and ABA implies that these seedlings retained at least part of the seed-maturation program, which is consistent with the continued expression of genes encoding oleosin and seed-storage proteins (see below).

Among the 232 genes commonly downregulated in the 3-day-old *drol1* seedlings when compared with the wild-type, those related to “mitotic cell cycle transition” and “regulation of cyclin-dependent protein serine/threonine kinase activity” were overrepresented (Table S4 and S5). Since many genes essential for the cell cycle progression and rapid vegetative development of seedlings were enriched in these GO terms, their downregulation in *drol1* seedlings is consistent with the abnormal growth of these seedlings.

### Derepression of seed-maturation genes after germination in *drol1* mutants

The *Arabidopsis* genome contains five genes (*OLE1–5*) that encode seed-specific oleosin proteins (D’Andréa *et al*., 2007; Miquel *et al*., 2014). These *OLE* genes exhibit high expression levels during seed maturation (Miquel *et al*., 2014) but are downregulated approximately 3 or 4 days after germination (DAG). The *drol1* mutant was originally identified based on the high level of *OLE3:LUC* expression in the seedlings 3 or 4 DAG (Suzuki *et al*., 2018). We surveyed the data for the mRNA levels of *OLE1–5* in the comprehensive transcriptomic analysis described above. In the wild-type seedlings, levels of all *OLE* mRNAs were very low at 3 DAG (Figure 5a–e). By contrast, significant levels of these *OLE* mRNAs were present in 3-day-old *drol1-1* mutant seedlings, and this rapidly decreased at 5 DAG. Similar patterns were observed in the *drol1-2* seedlings, although levels of the *OLE* mRNAs were lower than those in the *drol1-1* mutant. These changes in the levels of *OLE* mRNAs were similar to the expression of the *OLE3:LUC* reporter gene (Suzuki *et al*., 2018).

**Figure 5.**
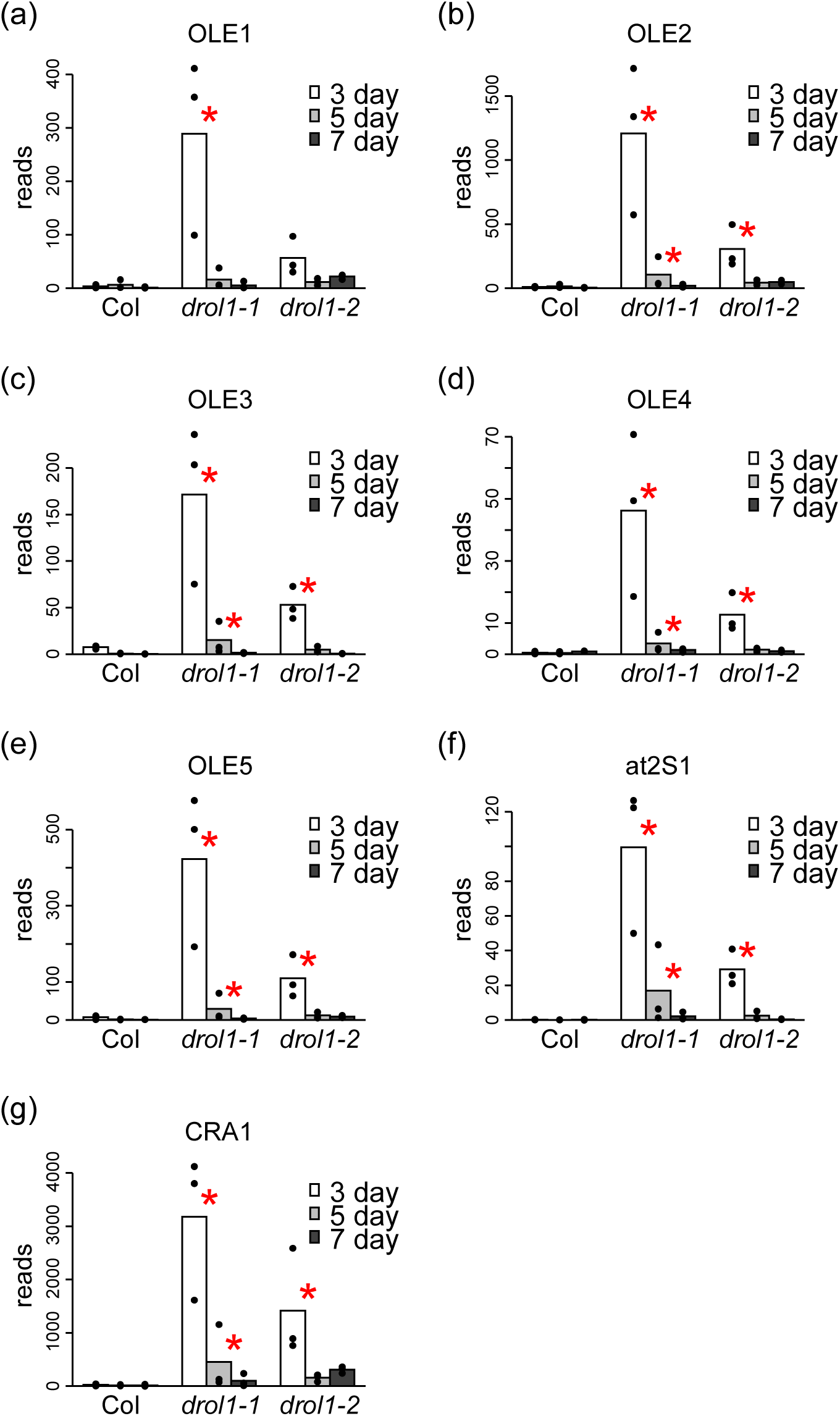
Differentially expressed genes (DEGs) in the *drol1* mutants. (a–e) Expression levels of five *OLEOSIN* (*OLE*) genes in the wild-type and *drol1-1* and *drol1-2* mutants. Short reads mapped to each gene were counted, and the counts were normalized against the total mapped reads. Bars and dots indicate mean (*n* = 3) and individual data, respectively. Asterisks indicate significantly increased expression compared with the wild-type at the same stage (FDR < 0.001). (f, g) Expression levels of 2S albumin (*at2S1*) and 18S globulin (*CRA1*), respectively.

The major seed-storage proteins in *Arabidopsis* are 2S albumin and 18S globulin (Fujiwara *et al*., 2002). Here we examined the effects of the *drol1* mutation on the expression of five 2S albumin genes and three 18S globulin genes after germination (Figure 5f and 5g and Figure S10). Similar to the *OLE* genes, significant levels of mRNA for these seed-storage protein genes were detected in the 3-day-old *drol1-1* seedlings when compared with the wild-type, and there was a rapid decline at 5 DAG. However, unlike the *OLE* genes, statistically significant differences in the levels of expression for the *drol1* mutants were observed only in genes with the highest levels of expression from each family. Only *at2S1* of the five 2S albumin and *CRA1* of the three 18S globulin genes were significantly upregulated in the *drol1* mutants (comparisons of Figure 5f and Figure S10a–d as well as Figure 5g and Figure S10e and f revealed this finding). The effects of the *drol1-2* mutation on the transcriptomic changes were considerably weaker than the *drol1-1* mutation. These results suggest that the defects in DROL1 caused the derepression of genes encoding not only oleosin proteins but also seed-storage proteins after germination.

### Induction of ABA responsible genes during germination in *drol1* mutants

The most important gene responses to ABA during germination were those of the bZIP transcription factor ABI5 and the B3 domain-containing transcription factor ABI3. The expression of both genes was induced by ABA, resulting in growth arrest during germination (Lopez-Molina *et al*., 2001). Both *ABI5* and *ABI3* showed significantly higher level of expression in *drol1* mutants compared with the wild-type (Figure S11a and b). The expression of *AtEm6* and *AtEm1*, which encode late embryogenesis abundant protein, is regulated by ABI5 downstream of ABI3 (Lopez□Molina *et al*., 2002). They were also upregulated in *drol1* mutants (Figure S11c and d). These results were consistent with the enrichment of the GO term “response to abscisic acid” in the upregulated genes in *drol1* mutants, and “mitotic cell cycle transition” in the downregulated genes in *drol1* mutants, when the activation of ABI5 and ABI3 causes growth arrest in the *drol1* mutants.

### Defective growth of *drol1* seedlings

In addition to the derepression of *OLE* genes, the *drol1* seedlings showed some developmental defects. The *drol1-1* and *drol1-2* mutant seedlings were smaller in size than the wild-type at 5-days (Figure 6a). While the wild-type cotyledons were curled upward, mutant cotyledons were either flat or occasionally curled downward. In addition, the root length of the *drol1* mutant seedlings was approximately 50% or 33% shorter than that of the wild-type seedlings at 3 and 5 DAG, respectively (Figure 6b; *p* < 0.5 with Tukey–Kramer test). However, root elongation between 5- and 7-day-old *drol1* seedlings was similar to that of the wild-type seedlings. Leaf development was also delayed in the *drol1* mutants when compared to the wild-type; when the wild-type seedlings possessed four leaves, the *drol1* mutant seedlings had fewer and smaller rosette leaves (Figure S2b). These growth defects were observed only when the *drol1* seedlings were grown on normal medium (as described in the Experimental procedure section). After planting in the soil, the growth defects were rescued, and the fertility of the *drol1* mutant seedlings was similar to that of the wild-type (Figure S12).

**Figure 6.**
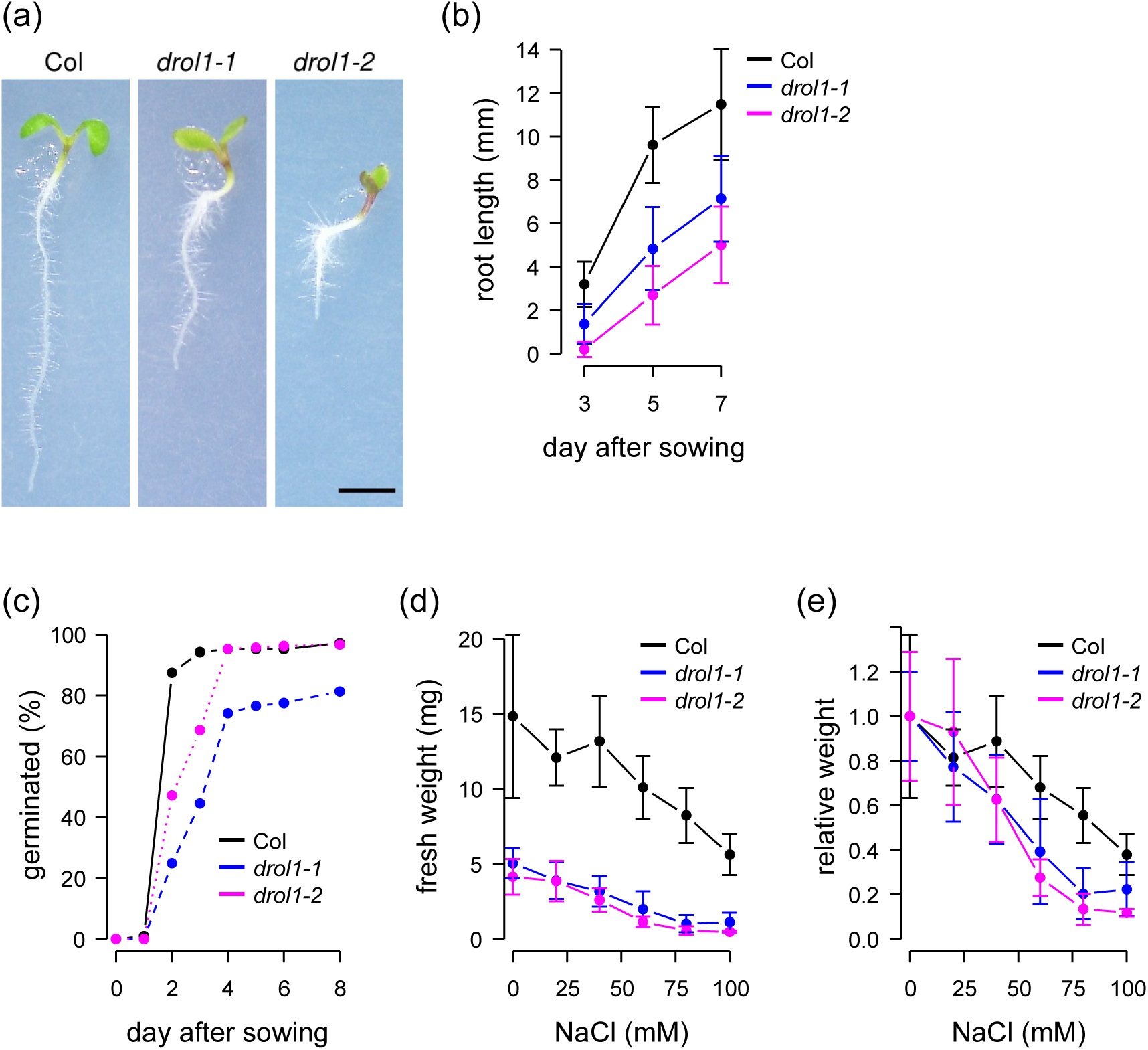
Phenotypes of the *drol1* mutant. (a) Photographs of 5-day-old wild-type (Col), *drol1-1*, and *drol1-2* seedlings (from left to right). Scale bar = 2 mm. (b) Root length of 3–7-day-old seedlings. Error bars represent standard deviation (*n* = 23–35). All differences between wild-type and *drol1* mutants at same days were statistically significant (*p* < 0.01) with Tukey–Kramer test. (c) Seed germination rate (*n* = 107–206). (d) Fresh weight of 10-day-old seedlings grown on medium supplemented with various concentrations of salt (NaCl) (*n* = 4–15). (e) Relative fresh weight for (d). The differences between wild-type and *drol1* mutants at 80 mM were statistically significant (*p* < 0.01) with Tukey–Kramer test.

As the genes involved in the response to dehydration and ABA were upregulated in the *drol1* seedlings, the phenotypes involving ABA were surveyed. The germination rate of the *drol1* mutant seeds was lower than that of the wild-type seeds (Figure 6c; *p* < 10^−90^ with chi-square test); all wild-type seeds germinated 2 days after sowing, whereas less than 50% of the *drol1* mutant seeds were germinated at this time, but the rate of germination gradually increased over a week. The germination process in the *drol1* mutants was delayed at lower concentration of ABA when compared with the wild-type. The openings of the cotyledons were inhibited on the medium containing 0.5 µM ABA in the *drol1* mutants and the greening of the cotyledons was also inhibited at 1 µM (Figure S13).

ABA has an important role in plant responses to high osmolarity and the genes for “response to salt stress” were overrepresented in the *drol1* mutants (Table S3) and the double mutant for *NHX5* and *NHX6*, encoding endosomal Na^+^/H^+^ antiporters, was reported to be sensitive to high osmolarity (Bassil *et al*., 2011). We also measured the fresh weight of 10-day-old wild-type, *drol1-1*, and *drol1-2* seedlings grown on medium containing various concentrations of salt (Figure 6d and e). The results indicated that *drol1* mutants were hypersensitive to salt (at 80 mM *p* < 0.01 with Tukey–Kramer test). Together, these results suggest that growth defects in the *drol1* mutants were at least partly caused by extended dormancy of the seeds and high osmolarity of the growth medium.

## Discussion

### DROL1 is a specific component of the spliceosome required for AT–AC-type intron splicing

In this study, we have shown that DROL1 is homologous to yeast DIB1, a highly conserved protein subunit of U5 snRNPs, but that it is also member of another subfamily specific to a certain class of eukaryotes (Figure 1). Defects in DROL1 specifically increased the retention levels of AT–AC-type introns (Figures 2 and 3 and Table 1).

Since *DROL1* is homologous to the *DIB1* gene of *S. cerevisiae* and the *hDim1* gene of *H. sapience*, it was hypothesized that DROL1 could mediate AT–AC-type intron splicing as a component of the U5 snRNP. Previous studies on *DLP/hDim2*, the human ortholog of *DROL1*, support this hypothesis, as it was shown to bind Prp6, a protein subunit of the U5 snRNP that interacts with DIB1 *in vitro* (Sun *et al*., 2004). Crystal structure analysis revealed that DLP/hDim2 shares a common overall structural fold with hDim1 (Simeoni *et al*., 2005). These results support the possibility that DLP/hDim2 replaced the position of hDim1 in the U5 snRNP to form a specific subclass of U5 snRNP.

The structural analysis of the human major spliceosome in the B complex revealed that hDim1 and Prp8, large scaffold proteins in the U5 snRNP, create a 5′ splice site binding pocket (Bertram *et al*., 2017). In this model, N98 and N99 of hDim1 contact the second 5′ terminal nucleotide of the intron. Indeed, our analysis of another cryo-EM structure of the human spliceosomal B complex reported recently (Zhan *et al*., 2018) revealed that residues of T96-G97-N98-N99 of hDim1 located very close to the 5′ end GU dinucleotide of the intron (Figure S14). These results suggest that the hDim1 protein plays a direct role in 5′ splice site recognition in the B complex (Bertram *et al*., 2017).

A precise structural comparison of hDim1 and DLP/hDim2 revealed two major differences, one of which is the additional β strand (referred to as β5 strand in the reference) specific to DLP/hDim2 (Simeoni *et al*., 2005). The β5 strand in DLP/hDim2 replaces a loop structure in hDim1 containing the conserved TGNNN amino acid sequence (underlined in Figure 1) that contacts with the 5′ end of the intron. The corresponding amino acid sequences of the DROL1 subfamily are T/SPDHT and are quite different from those of TGNNN in the DIB1 subfamily. This structural difference between hDim1 and DLP/hDim2 is possibly associated with the difference in the 5′ splice site sequence.

Schreib *et al*. (2018) introduced missense mutations into the yeast DIB1 gene to create temperature-sensitive mutants (Schreib *et al*., 2018). Spliceosome assembly was analyzed in these mutants and it was found that the stepwise formation of the spliceosome was stalled at the B complex under nonpermissive temperature. If this was applied to the *drol1* mutant, then the AT–AC-type-specific spliceosome without functional DROL1 would also stall on the AT–AC intron. This might protect premature mRNA from degradation by nonsense mediated decay and enable detection of intron retention in the *drol1* mutant.

The results presented in this study strongly suggest the presence of AT–AC-type-specific spliceosomes containing DROL1; however, no difference in protein component between the U4_atac_/U6_atac_.U5 tri-snRNPs and U4/U6.U5 tri-snRNPs was reported (Schneider *et al*., 2002). Immunoprecipitation experiments were conducted with antibodies against proteins of the major tri-snRNP and they succeeded in detecting U4_atac_ and U6_atac_ snRNAs in precipitants. From these results it was concluded that the protein compositions of U4_atac_/U6_atac_.U5 tri-snRNPs were very similar to U4/U6.U5 tri-snRNPs, if not identical. However, as immunoprecipitation with antibodies against hDim1 and DLP/hDim2 has not been tested, it is not clear whether DLP/hDim2 is present in U4_atac_/U6_atac_.U5 tri-snRNPs.

Recently, cryo-EM structures of the human minor spliceosomal C complex were reported (Bai *et al*., 2021). However, hDim2/DLP was not identified in the complex. Considering the disassembly of hDim1 from the spliceosome before the transition to the major C complex, the absence of hDim2/DLP from the minor C complex would be reasonable. Identification of minor spliceosomal B complex containing DROL1 subfamily and proteins interacting with the DROL1 subfamily are needed to elucidate how DROL1 mediates AT–AC intron splicing.

### *drol1* phenotypes dependent on ABA likely result from defective AT–AC-type intron splicing

The RNA-Seq analysis showed that many genes were upregulated or downregulated in *drol1* when compared with the wild-type (Tables S2–S5 and Figure S9). All five seed-specific *OLE* genes, one of which was used as a reporter for the screening of the *drol1-1* mutant, were upregulated in the *drol1* mutants (Figure 5), confirming our previous results (Suzuki *et al*., 2018). These transcriptomic changes were speculated to cause pleiotropic phenotypes of the *drol1* mutant such as expression of seed-maturation genes in seedlings, delayed seedling growth, low seed germination rate, and hypersensitivity to salt (Figure 6).

The most prominent changes in the *drol1* mutants at the transcription level included the upregulation of several ABA-responsive genes, including *ABA INSENSITIVE 5* (*ABI5*) (Figure S11). *ABI5* expression is known to be induced by the application of ABA to germinating embryos, resulting in growth arrest during germination (Lopez-Molina *et al*., 2001). The ABI5 protein induces the expression of *AtEm1* and *AtEm6*, which are ordinarily expressed during late embryogenesis (Lopez-Molina *et al*., 2002), and these inductions also occurred in *drol1* seedlings (Table S2 and Figure S11). Consistent with the retarded growth, the genes related to mitosis were downregulated in *drol1* seedlings (Tables S4 and S5). Based on these results, we hypothesized that the activation of ABA signaling in the *drol1* seedlings, which in turn activated *ABI5* expression, resulted in the retardation of germination and seedling growth.

The activation of ABA signaling in the *drol1* mutants is speculated to be caused by pre-mRNAs of genes harboring AT–AC-type introns. Two candidate genes accountable for the activation of ABA signaling, *NHX5* and *NHX6*, encode endosomal Na^+^/H^+^ antiporters (Table 1). Both *NHX5* and *NHX6* uniquely contain two AT–AC-type introns, which are conserved in human orthologous genes, *NHE6* and *NHE7* (Zhu and Brendel, 2003). The *NHX5* and *NHX6* genes are involved in salt tolerance in plants, and the *nhx5 nhx6* double mutant plants are smaller than wild-type (Bassil *et al*., 2011). When two AT–AC-type introns were independently retained in the transcriptome of *drol1*, the level of mature *NHX5* and *NHX6* mRNAs was expected to be severely reduced, thus mimicking the hypersensitivity of the *nhx5 nhx6* double mutant to salt. This hypersensitivity might cause the induction of the genes related to the “response to salt stress” and activation of ABA signaling in *drol1* mutants.

The *HD2* gene family is another candidate that could account for the *drol1* mutant phenotypes. The *HD2* gene family encodes plant-specific histone deacetylases, which remove the acetyl group from histones H3 and H4 and generally repress gene expression (Hollender and Liu, 2008). The *Arabidopsis* genome harbors four *HD2* genes, and in three of these genes (except *HD2A*), the third intron is an AT–AC type. The *hd2c* mutant exhibits lower germination than the wild-type on medium containing ABA and NaCl (Colville *et al*., 2011; Luo *et al*., 2012). Furthermore, as plants overexpressing *HD2D* showed tolerance to NaCl, HD2D is suggested to play a role in the responses to high salinity conditions (Han *et al*., 2016). It was difficult to attribute the pleiotropic phenotype of the *drol1* mutant to HD2C alone due to the normal development of the *hd2c* mutants in the absence of abiotic stress. In our experiments, the retarded growth of the *drol1-2* mutant was not complemented by the modified *HD2B* or *HD2C*, wherein the AT–AC-type introns were substituted to GT–AG-type introns. Moreover, the modified *HD2D* or *NHX5* genes, wherein AT–AC-type introns were deleted, also failed to complement the *drol1* mutants (Figure S15). However, it is possible that the reduced expression of both *HD2C* and *HD2D* increased the sensitivity to ABA and salt stress. It is also possible that the reduction in *HD2C* and *HD2D* expression enhanced the sensitivity to ABA and salt due to *NHX5* and *NHX6* defects.

### Survival of *drol1* mutants with critical genes that have AT–AC-type introns

Some of the genes essential for cell viability also have AT–AC-type introns. Homozygous mutants of the *NRPA2* gene, which encodes the second largest subunit of the nuclear RNA polymerase I, were lethal (Onodera *et al*., 2008). The first intron of *NRPA2* was the AT–AC type, and only 10% of the transcripts were spliced (Figure S7). A small amount of the spliced *NRPA2* transcript might enable reduced growth. Moreover, the insertion of polypeptides translated from the retained 96-bps first intron and no stop codon might not disrupt the function of RNA polymerase I. *DPB2* encodes the regulatory subunit of DNA polymerase ε, and homozygous *dpb2* mutants are lethal (Ronceret *et al*., 2005). The *SMC1* gene, which encodes a subunit of cohesin, is essential for embryogenesis (Liu *et al*., 2002). These results indicate that a part of the pre-mRNA with an AT–AC-type intron was spliced in the absence of a functional DROL1.

Recently, it was reported that a part of human mosaic variegated aneuploidy was caused by a mutation in *CENATAC*, which caused the defects in splicing of AT–AC-type introns (de Wolf *et al*., 2021). Evolutionary co-occurrence analysis revealed that *CENATAC* had the strongest linkage with DLP/hDim2 (referred to as TXNL4B in the reference), a human ortholog of *DROL1*. The interactions between CENATAC and DLP/hDim2 were also confirmed by co-immunoprecipitation (de Wolf *et al*., 2021). The ortholog of CENATAC in *Arabidopsis* is *TITAN-LIKE* (*TTL*), and the *ttl* mutant was screened for the defective development of embryo and endosperm (Lu *et al*., 2012). Three of the original nine *titan* (*ttn*) loci, consisting of *ttn8*, *ttn3*, and *ttn7*, encoded subunits for the structural maintenance of chromosomes, SMC1, SMC2, and SMC3, respectively (Liu *et al*., 2002). Although the defective splicing of AT–AC-type introns in SMC1 mRNA, which occurred in *drol1* mutants, has not been reported in *ttl* mutants, it is speculated that the similar phenotypes of the *ttl* and *ttn8* mutants are caused by defects in SMC1. Because both *CENATAC* and *TTL* have AT–AC-type introns and splicing disrupts their function, the activity of AT–AC-type-specific spliceosome may be regulated in a feedback manner (de Wolf *et al*., 2021). Because the splicing of the AT–AC-type intron in *TTL* was almost completely retained in *drol1* mutants (PIR = 100 in *drol1-1* and 97 in *drol1-2*), it was speculated that the amount of TTL protein was increased. This might activate the AT–AC-type-specific spliceosome and save *drol1* mutants from lethality.

Differences between the transcriptomes of wild-type and *drol1* seedlings, in terms of both gene expression and intron retention, gradually decreased with increasing age (Figure S9). Consistent with these results, *drol1* developmental defects were rescued by planting the seedlings in soil (Figure S12). These results also support the idea that DROL1 is required, but not essential, for the effective splicing of AT–AC-type introns. The time-dependent accumulation of mature mRNAs might help rescue the lethal phenotype of *drol1* mutants.

## Experimental procedures

### Comparison of *DROL1* homologs

Amino acid sequences of the DROL1 homologs from different organisms were downloaded from different databases: *A. thaliana* from the TAIR database (https://www.arabidopsis.org/), *Oryza sativa* from the Rice Annotation Project Database (RAP-DB; http://rapdb.dna.affrc.go.jp/), *Physcomitrella patens* and *Chlamydomonas reinhardtii* from the Phytozome database (https://phytozome-next.jgi.doe.gov/), *Caenorhabditis elegans* from the WormBase (https://www.wormbase.org/), *Schizosaccharomyces pombe* from the PomBase (https://www.pombase.org/), and *Saccharomyces cerevisiae* from the *Saccharomyces* Genome Database (https://www.yeastgenome.org/). *Mus musculus* and *Homo sapiens* homologs of DROL1 were found in the UniProt database through the DNA Data Bank of Japan (DDBJ) blastp server. Since the *C. reinhardtii* homolog of DROL1 was truncated at the N-terminus, full-length sequences were searched using exonerate (Slater and Birney, 2005), with AtDIB1 (AT5G08290.1) as the query. Multiple sequence alignments and a phylogenetic tree were constructed using ClustalW2 (Larkin *et al*., 2007) and modified manually.

### Plant material and growth conditions

*A. thaliana* (L.) Heynh. strain Columbia (Col-0) was used as the wild-type. *drol1-1* and *drol1-2* were described previously (Suzuki *et al*., 2018). Unless otherwise indicated, seeds were sterilized on 0.3% (wt/vol) gellan gum plates containing half strength Murashige and Skoog medium (Wako, Japan) supplemented with 0.05% (wt/vol) 2-morpholinoethanesulfonic (MES) acid (pH 5.7), 100 mg/L myo-inositol, 10 g/L thiamine HCL, and 2% (wt/vol) sucrose. Plates were wrapped with aluminum foil and incubated at 4°C for 2–4 days. Plates were incubated in a growth chamber at 22°C under continuous fluorescent light at an intensity of 65 µmol m^-2^ s^-1^.

### RNA extraction

Sterilized seeds were sown on 1.5% (w/v) agar plates containing medium. After vernalization, plates were vertically placed in a growth chamber at 22°C under continuous fluorescent light. Seedlings were collected after 3, 5, and 7 days of incubation, immediately frozen in liquid nitrogen, and ground to a fine powder using a pestle and mortar. Total RNA was extracted from the powder using the NucleoSpin RNA Plant Kit (Takara; http://www.takara-bio.co.jp/), according to the manufacturer’s protocol.

### 5′ RACE analysis of the *DROL1* transcript

The 5′ RACE analysis was conducted using the 5′ -Full RACE Core Set (Takara), according to the manufacturer’s protocol. The cDNAs were reverse transcribed using a 5′ -phosphorylated primer 5RACE_RT purchased from Eurofins Genomics (https://www.eurofinsgenomics.jp/; all sequences of primers were shown in Table S6). The first and second amplifications of cDNA were performed using the following primer pairs: 5RACE_S1 and 5RACE_A1 for the first amplification and 5RACE_S2 and 5RACE_A2 for the second amplification. The PCR products were purified and ligated to adapters for NextSeq500 (Illumina, http://www.illuminakk.co.jp/). A total of 451,660 reads were obtained, of which 197,822 (44%) contained a reverse-transcription primer sequence. After trimming the primer sequence, reads were aligned to the reference sequence that contained a 2-kbp promoter and *DROL1* coding sequence using Bowtie (Langmead *et al*., 2009). The aligned 160,418 reads (81% of trimmed reads) were counted at each position at which they originated.

### RNA-Seq analysis

The extracted total RNA was used to construct cDNA libraries with the NEBNext Ultra RNA Library Prep Kit for Illumina, NEBNext Poly(A) mRNA Magnetic Isolation Module, and NEBNext Multiplex Oligos for Illumina (New England BioLabs, https://www.nebj.jp/), according to the manufacturer’s protocol. The libraries were sequenced using the NextSeq500 sequencer (Illumina). Raw reads containing adapter sequences were trimmed using bcl2fastq (Illumina), and nucleotides with low-quality (QV < 25) were masked by N using the original script. Reads <50 bps were discarded, and the remaining reads were mapped to the cDNA reference sequence using Bowtie with the following parameters: “--all --best --strata” (Langmead *et al*., 2009). Reads were then counted by transcript models. DEGs were selected based on the adjusted *p*-value calculated using edgeR (version 3.20.9) with default settings (Robinson *et al*., 2010).

### Analysis of the retained introns

Reads obtained by RNA-Seq analysis were mapped to the genomic sequence using Tophat (Trapnell *et al*., 2009) with default parameters. PIR, logFC, and FDR for the introns were calculated using the anchorbased function of ASpli (Estefania *et al*., 2021).

### Gene cloning and plasmid construction

DNA fragments for the *DROL1*, *HD2B*, and *HD2C* genes were amplified with specific primers (Table S6) and cloned into the pDONR vector using Gateway BP clonase (Thermo Fisher Scientific, https://www.thermofisher.com/). The nucleotide substitutions from AT–AC to GT–AG in *HD2C* were introduced using the QuickChange Multi Site-Directed Mutagenesis Kit according to the manufacturer’s protocol (Agilent, https://www.chem-agilent.com/). The plasmids for the *HD2B* and *DROL1* genes with nucleotide substitutions and insertions were obtained by inverse-PCR and self-ligation with specific primers. The inserts were transferred into pGWB501 binary vector (Nakagawa *et al*., 2007), and then used for transformation.

### RT-PCR

cDNA was reverse transcribed from 1 µg of total RNA by SuperScript III (Thermo Fisher Scientific) with 2 pmol of each of the five anti-sense primers. The synthesized cDNA was amplified with PCR using TaKaRa Ex Taq (Takara) over 35 cycles and applied for chip electrophoresis using MultiNA (Shimadzu, https://www.an.shimadzu.co.jp/). All primers used have been listed in Table S6.

### Physiological experiments

To measure the root length, seedlings were grown on medium solidified with 1.5% agar and the plates were kept vertical. To measure the germination rate, seeds were placed on the normal medium and the tearing of the seed coat was counted as germination under stereomicroscope (1–4×). For the salt tolerance test, seeds were placed on normal medium containing 0, 20, 40, 60, 80, and 100 mM NaCl and grown for 10 days in a growth chamber after 2-days of vernalization.

### Statistical analysis

All statistical tests in this study were performed on R (https://www.r-project.org/). For Student’s *t*, chi-square, and Tukey–Kramer, t.test, chisq.test, and TukeyHSD functions were used, respectively.

### Accession number

Sequence data from this article can be found in the DDBJ Sequence Read Archive under the accession number DRA010788.

## Acknowledgments

We thank Sumie Ishiguro and Hironaka Tsukagoshi for critically reading our work. We also thank Ayami Furuta for technical assistance. This work was supported in part by Grant-in-Aid from the Ministry of Education, Culture, Sport, Science and Technology (17K07454 to T.S.), from Chubu University (19M04A1 to T.S.), and by the Japan Science and Technology Agency (ERATO JPMJER1004 to T.H.). The authors would like to thank Enago (www.enago.jp) for English language review.

---

Table S1. Summary of the RNA-Seq analysis performed in this study

Table S2. Upregulated genes in drol1 seedlings 3 days after germination (DAG)

Table S3. Gene ontology (GO) analysis of the upregulated genes in the *drol1* mutants

Table S4. Downregulated genes in drol1 seedlings 3 DAG

Table S5. GO analysis of the downregulated genes in the *drol1* mutants

Table S6. Primers used in this study

Figure S1. Multiple sequence alignment for the DROL1 homologs

Figure S2. Analysis of the transcription and translation start site of *DROL1*

Figure S3. Scheme for the analysis of retained and spliced introns

Figure S4. Analysis of intron retention in the *drol1-2* mutants

Figure S5. Retention of the third intron in *HD2C* mRNA

Figure S6. Retention of the third intron in *HD2D* mRNA

Figure S7. Retention of the first intron in *NRPA2* mRNA

Figure S8. Retention of the third and eleventh introns in *NHX5* mRNA

Figure S9. Differentially expressed genes (DEGs) in *drol1* mutants

Figure S10. Increased expression of genes encoding seed-storage proteins in *drol1* seedlings

Figure S11. Increased expression of genes responsible to ABA in *drol1* seedlings

Figure S12. Plant phenotypes of *drol1-1* mutants grown on soil

Figure S13. Seedlings germinated on medium containing ABA

Figure S14. Interaction between hDim1 and pre-mRNA

Figure S15. Photographs of the *drol1* mutants transformed with intron-modified genes

