## Supplementary material for "*DROL1* subunit of U5 snRNP in the spliceosome is specifically required to splice AT–AC-type introns in *Arabidopsis*": Table 1

Table 1. Genes with AT-AC-type introns showing significant intron retention in *drol1* mutants

| Gene | dPIR | logFC | FDR | Gene function |
| --- | --- | --- | --- | --- |
| AT5G22650 | 86.3 | -2.65 | 9.33e-107 | histone deacetylase 2B / HD2B |
| AT4G13615 | 31.9 | -0.66 | 1.81e-57 | Uncharacterised protein family SERF |
| AT3G54670 | 90.5 | -3.26 | 4.06e-52 | Structural maintenance of chromosomes family protein / SMC1 |
| AT1G29940 | 90.1 | -3.13 | 1.86e-49 | nuclear RNA polymerase A2 / NRPA2 |
| AT4G15802 | 17.3 | -0.35 | 3.01e-44 | heat shock factor binding protein |
| AT5G56900 | 74.4 | -1.94 | 5.06e-44 | CwfJ-like family protein / zinc finger (CCCH-type) family protein |
| AT5G22110 | 87.7 | -2.98 | 1.46e-40 | DNA polymerase epsilon subunit B2 / DPB2 |
| AT1G80500 | 44.2 | -0.97 | 1.45e-39 | SNARE-like superfamily protein SNARE-like superfamily protein |
| AT3G62830 | 52.4 | -1.17 | 1.26e-38 | UDP-glucuronic acid decarboxylase 2 / UXS2 |
| AT3G53180 | 71.0 | -1.82 | 4.32e-38 | glutamate-ammonia ligases |
| AT3G51460 | 32.5 | -0.68 | 1.97e-34 | Phosphoinositide phosphatase family protein |
| AT5G18410 | 77.0 | -2.08 | 8.07e-32 | transcription activators |
| AT2G39080 | 24.8 | -0.51 | 3.24e-31 | NAD(P)-binding Rossmann-fold superfamily protein |
| AT3G22880^†2^ | 65.7 | -1.59 | 4.15e-28 | DNA repair (Rad51) family protein / DMC1 |
| AT3G50590 | 44.3 | -0.97 | 1.02e-27 | Transducin/WD40 repeat-like superfamily protein |
| AT4G38240 | 52.9 | -1.20 | 2.90e-27 | α-1,3-mannosyl-glycoprotein β-1,2-N-acetylglucosaminyltransferase, putative |
| AT1G06720 | 37.1 | -0.80 | 4.89e-27 | P-loop containing nucleoside triphosphate hydrolases superfamily protein |
| AT5G38380 | 55.7 | -1.29 | 1.97e-26 | unknown protein |
| AT3G06820 | 69.1 | -1.83 | 3.37e-24 | Mov34/MPN/PAD-1 family protein |
| AT2G40840 | 16.9 | -0.34 | 3.01e-23 | disproportionating enzyme 2 / DPE2 |
| AT1G29630 | 73.7 | -1.98 | 5.31e-23 | 5'-3' exonuclease family protein /EXO1 |
| AT5G27380 | 35.5 | -0.75 | 5.95e-23 | glutathione synthetase 2 / GSH2 |
| AT2G27840 | 18.1 | -0.37 | 4.92e-22 | histone deacetylase 2D / HD2D |
| AT1G77320 | 68.9 | -1.89 | 3.98e-21 | transcription coactivators |
| AT5G08430 | 71.5 | -2.22 | 1.73e-20 | SWIB/MDM2 and Plus-3 and GYF domain-containing protein |
| AT5G63700 | 84.4 | -3.25 | 2.82e-20 | zinc ion binding / DNA binding protein |
| AT3G21215 | 41.0 | -0.88 | 2.96e-20 | RNA-binding (RRM/RBD/RNP motifs) family protein |
| AT1G67960 | 60.8 | -1.50 | 5.48e-20 | unknown protein |
| AT1G73350 | 49.8 | -1.25 | 6.19e-19 | unknown protein |
| AT4G30900 | 65.3 | -1.63 | 7.08e-19 | DNAse I-like superfamily protein |
| AT1G80210 | 60.2 | -1.42 | 7.89e-19 | Mov34/MPN/PAD-1 family protein / BRCC36A |
| AT2G47650 | 18.1 | -0.37 | 2.50e-18 | UDP-glucuronic acid decarboxylase 4 / UXS4 |
| AT1G24050 | 10.6 | -0.21 | 8.31e-18 | RNA-processing, Lsm domain |
| AT5G03740 | 16.5 | -0.33 | 8.55e-18 | histone deacetylase 2C / HD2C |
| AT1G26170^†1^ | 47.9 | -1.06 | 1.56e-17 | ARM repeat superfamily protein |
| AT1G79610 | 56.0 | -1.31 | 2.40e-16 | Na^+^/H^+^ exchanger 6 / NHX6 |
| AT3G53520 | 14.6 | -0.30 | 1.78e-15 | UDP-glucuronic acid decarboxylase 1 / UXS1 |
| AT3G51120 | 48.3 | -1.08 | 5.33e-15 | zinc finger CCCH domain-containing protein 44 |
| AT3G24100 | 6.8 | -0.13 | 9.08e-15 | Uncharacterised protein family SERF |
| AT4G24900 | 54.7 | -5.77 | 8.24e-14 | C2H2-domain protein / TTL |
| AT1G31660 | 24.3 | -0.51 | 8.48e-14 | unknown protein |
| AT2G39960 | 8.8 | -0.18 | 3.40e-13 | Microsomal signal peptidase 26 kDa subunit / SPC25 |
| AT2G16485 | 17.1 | -0.35 | 2.84e-9 | NEEDED FOR RDR2-INDEPENDENT DNA METHYLATION / NERD |
| AT2G44270^†1^ | 20.0 | -0.40 | 2.90e-9 | repressor of *lrx1* / ROL5 |
| AT1G54370 | 31.8 | -0.66 | 5.61e-7 | Na^+^/H^+^ exchanger 5 / NHX5 |
| AT5G44200 | 13.0 | -0.27 | 5.94e-7 | CAP-binding protein 20 / CBP20 |
| AT1G76170 | 46.1 | -1.08 | 1.44e-5 | 2-thiocytidine tRNA biosynthesis protein / TtcA |
| AT1G02750^†1^ | 13.5 | -0.28 | 9.43e-5 | Drought-responsive family protein Drought-responsive family protein |

Genes are listed ascending to FDR.
Genes marked by †1 and †2 were only listed in the comparison to *drol1-1* and *drol1-2*, respectively.
