## Supplementary Figures for "*DROL1* subunit of U5 snRNP in the spliceosome is specifically required to splice AT–AC-type introns in *Arabidopsis*"

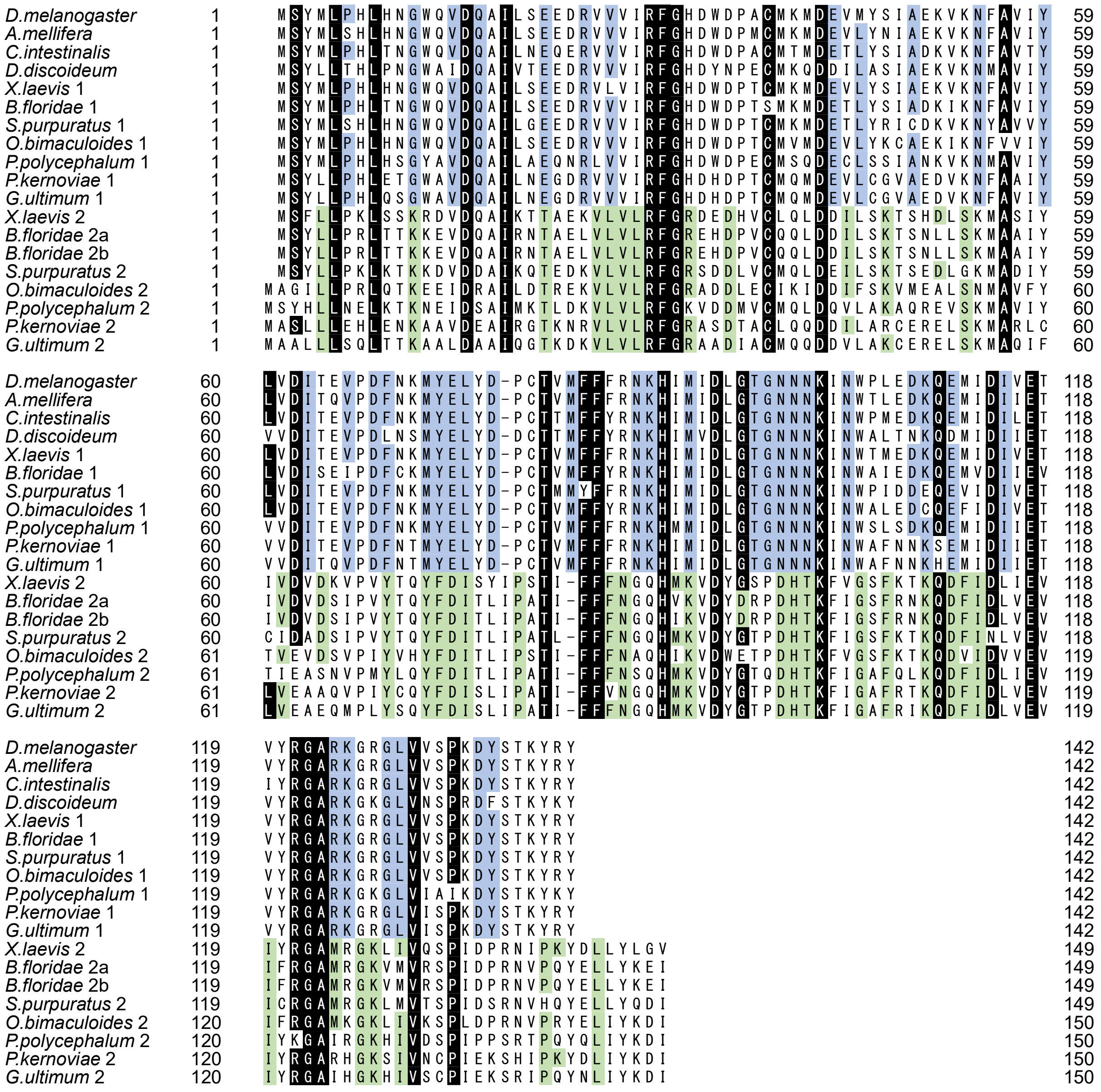


**Figure S1. Multiple sequence alignment for the DROL1 homologs**

Sources of the DROL1 amino acid sequences were as follows: *Drosophila melanogaster* (Q8SZ87_DROME in UniProt; https://www.uniprot.org/), *Xenopus laevis* (rna22650 rna37839 in Xlaevis 9.2; http://ftp.xenbase.org), *Apis mellifera* (genome assemblies from ftp:// ftp.ensemblgenomes.org), *Ciona intestinalis* and *Branchiostoma floridae* (genome assemblies from https://genome.jgi.doe.gov), *Strongylocentrotus purpuratus*, *Physarum polycephalum* , *Phytophthora kernoviae* , and *Globisporangium ultimum* : genome assemblies GCF_000002235.4, GCA_000413255.3, GCA_008080845.1, and GCA_000143045.1, respectively, from https://www.ncbi.nlm.nih.gov/. Open reading frames were identified in genome assemblies using tblastn and exonerate. Short reads of *Octopus bimaculoides* and *Dictyostelium discoideum* from the Short-Read Archive database (https://www.ncbi.nlm.nih.gov/sra) were assembled by Trinity. The rules for naming and coloring are consistent with those used in Figure 1.

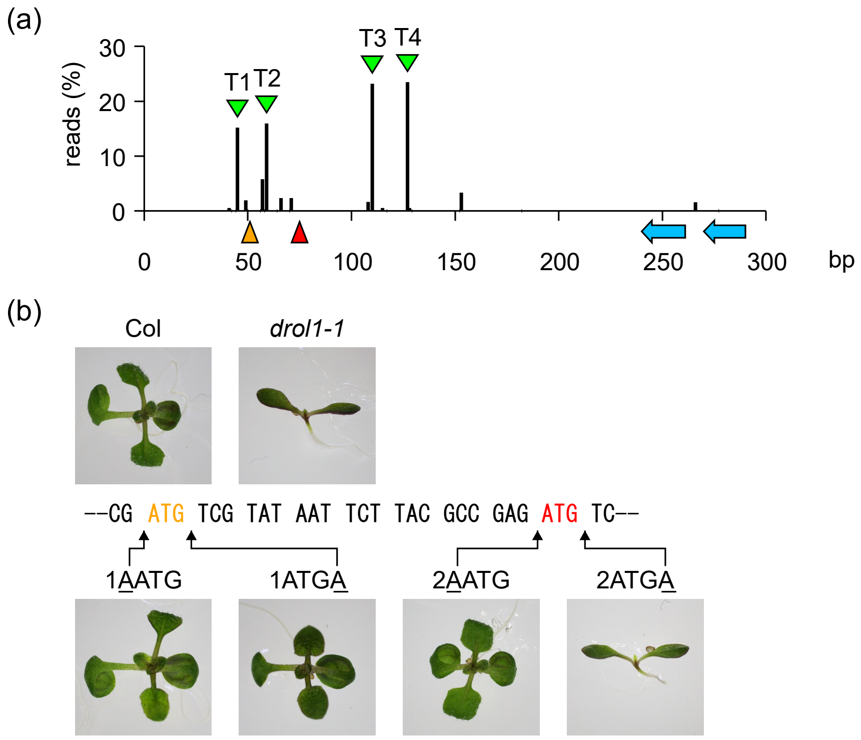


**Figure S2. Analysis of the transcription and translation start site of *DROL1***

(a) Transcription start sites determined by 5′ RACE. The 5′ ends of the aligned reads obtained by 5′ RACE were counted and plotted. Orange and red arrowheads indicate the putative translational start sites. Four major peaks of 5′ ends are indicated by green triangles. The blue arrows indicate 5′ RACE primers. (b) Analysis of the translational start site. Nucleotide sequences corresponding to the longer N-terminal amino acid sequences of DROL1 and two ATG codons (orange and red) are indicated. The insertion sites of adenine (A) nucleotide in four mutant variants of the *DROL1* gene (1AATG, 1ATGA, 2AATG, and 2ATGA) are indicated by arrows. Images of 10-day-old seedlings of the wild-type (Col) and *drol1-1* mutant and four transformants are shown. More than three independent lines were tested for each mutated gene.

(a)

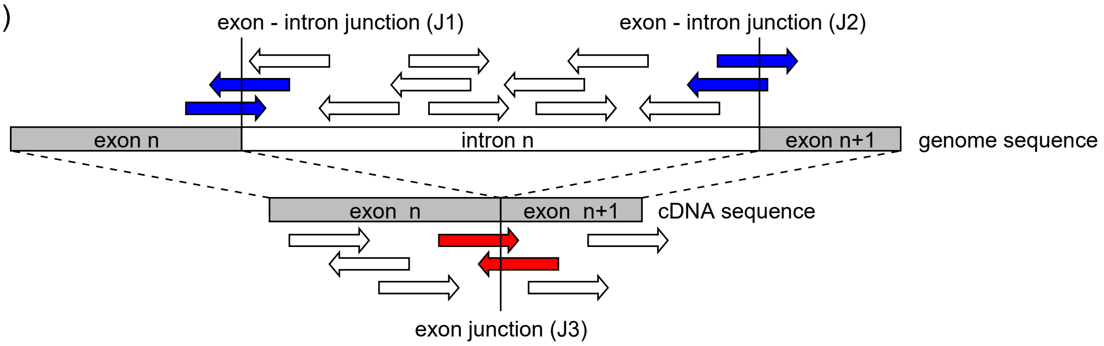


(b)

$$PIR = \frac{J1 + J2}{J3 \times 2 + J1 + J2} \times 100$$

**Figure S3. Scheme for the analysis of retained and spliced introns**

Short reads obtained by RNA-Seq were mapped to the genome reference. Reads spanning the exon–exon junctions (red arrows) and the exon–intron junction (blue arrows) were counted to determine the percent of intron retention (PIR) for each intron.

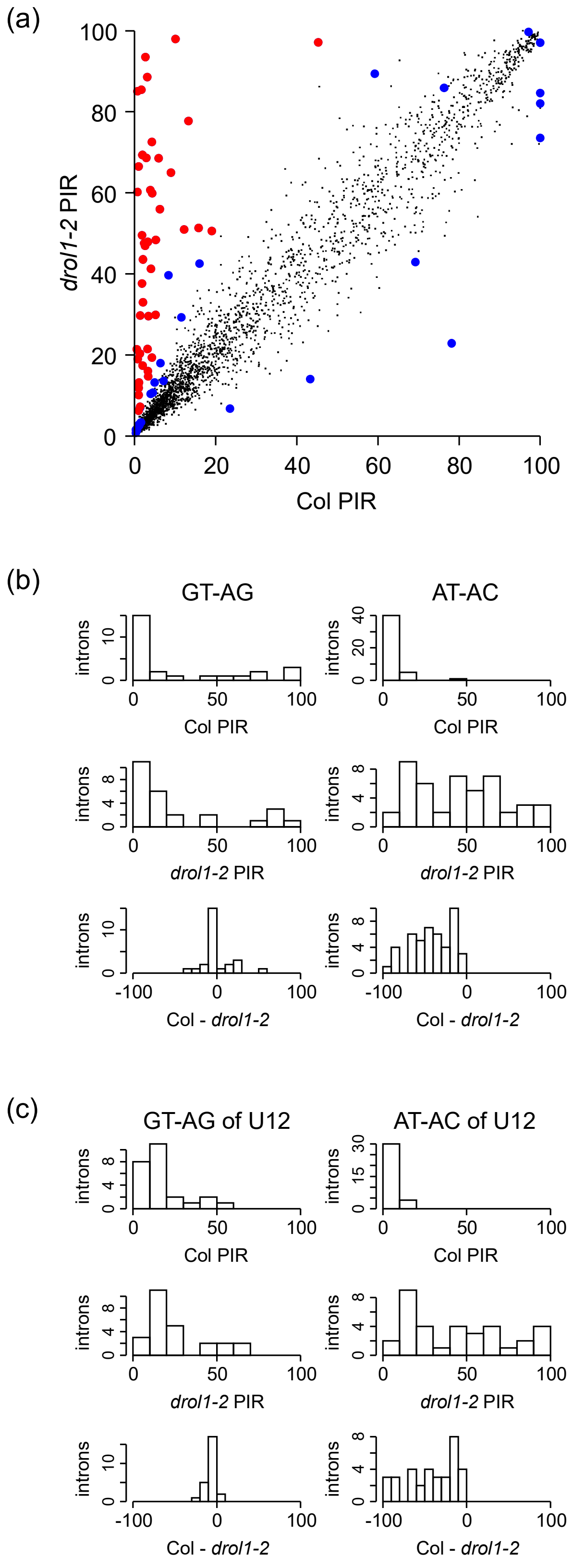


**Figure S4. Analysis of intron retention in the *drol1-2* mutants**

(a) Scatter plot of percent intron retention (PIR) for the wild-type (Col) and *drol1-2* mutant. Out of 2,930 introns analyzed, the splicing patterns of 75 (blue and read large dots) were significantly changed (FDR < 0.01). Red dots indicate 46 AT–AC-type introns, as in Figure 2a. (b) Histograms of PIR for 29 GT–AG- and 46 AT–AC-type introns. The first and second rows represent the observed PIR in the wild-type and *drol1-2*, respectively. The third row represents the differences in PIR between the wild-type and *drol1-2*. (c) Histograms of PIR for 25 GT–AG- and 34 AT–AC-type of U12-dependent introns.

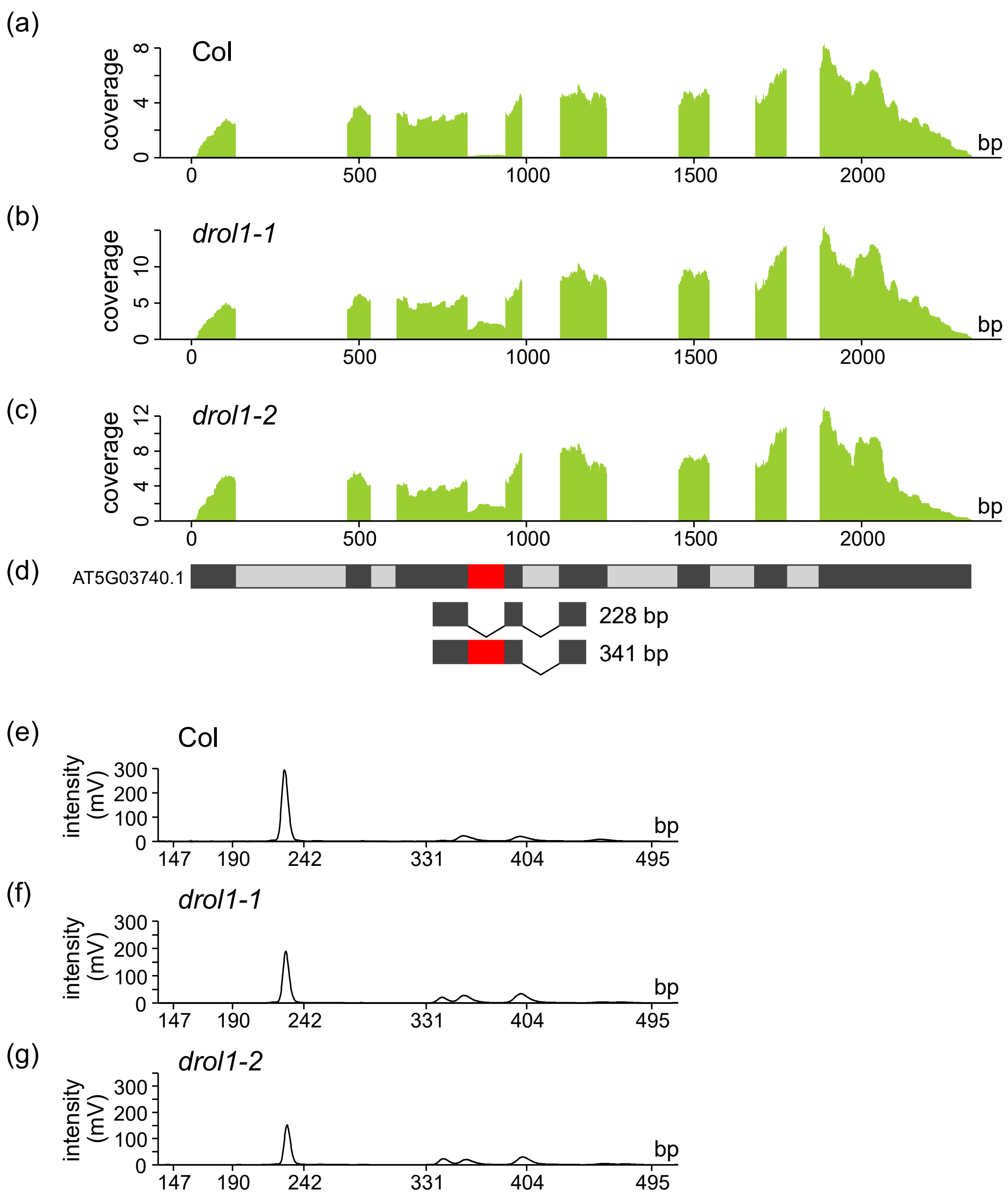


**Figure S5. Retention of the third intron in *HD2C* mRNA**

Analysis of intron retention in *HD2C* as indicated in Figure 3 for *HD2B*.

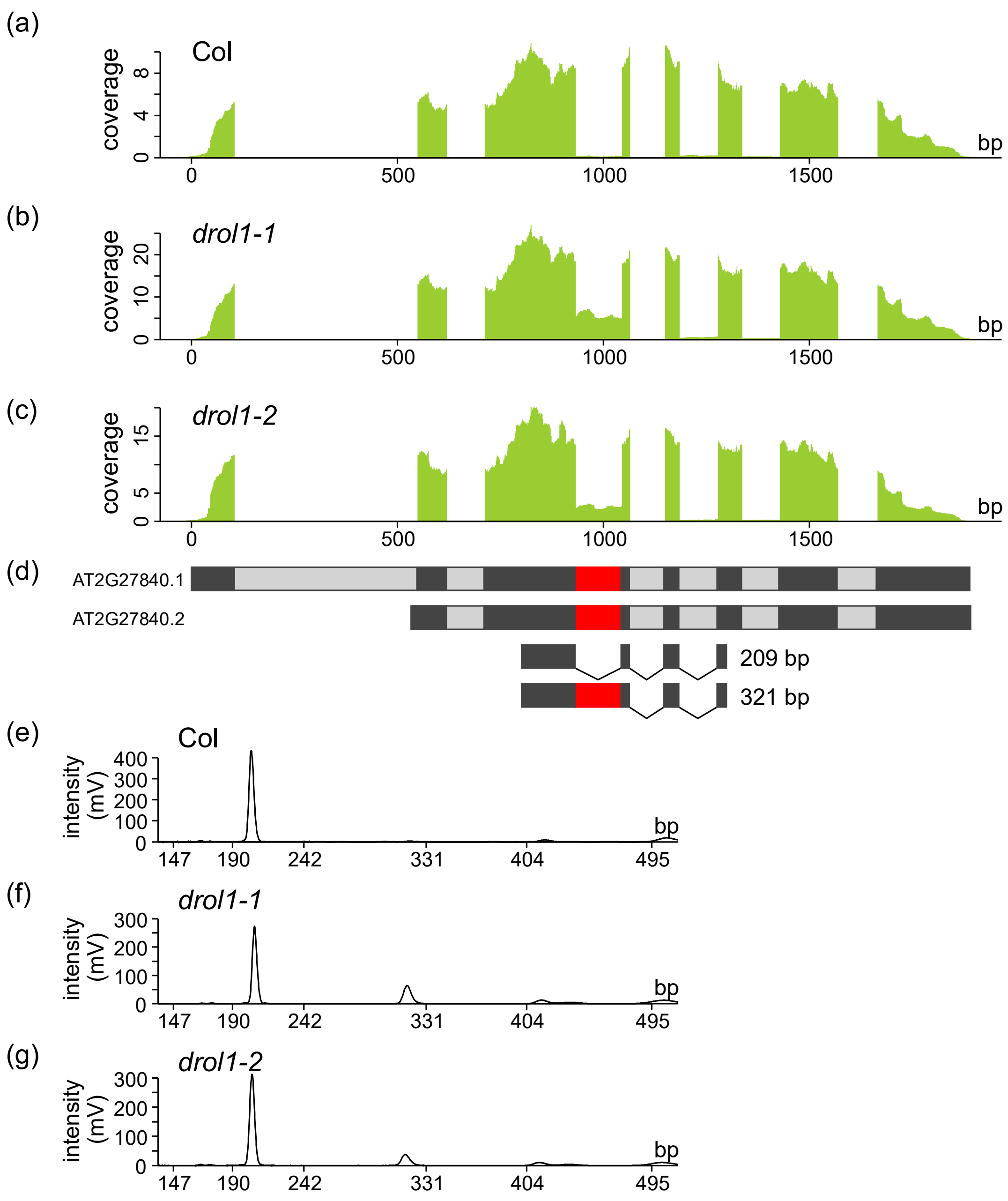


**Figure S6. Retention of the third intron in *HD2D* mRNA**

Analysis of intron retention in *HD2D* as indicated in Figure 3 for *HD2B*.

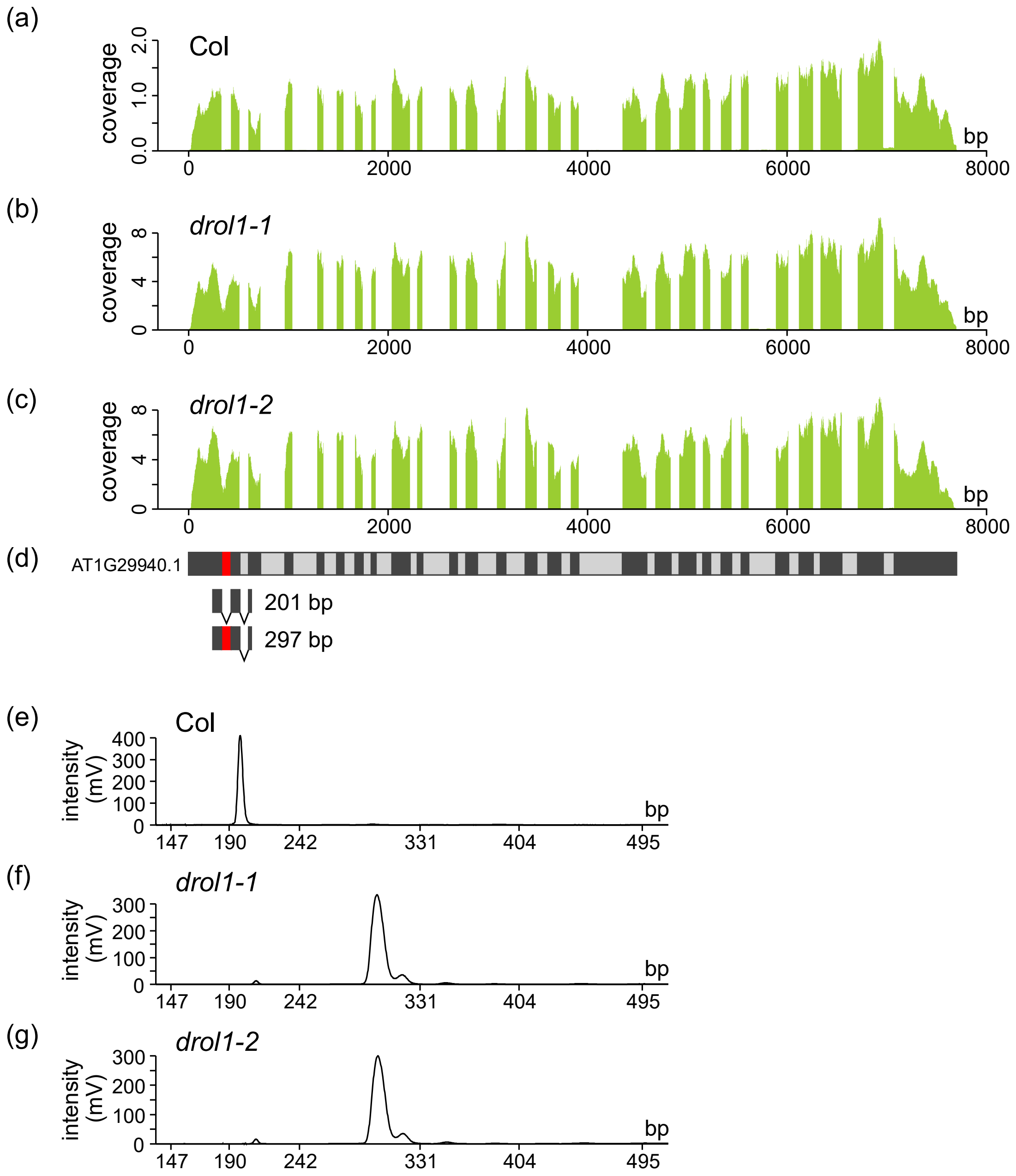


**Figure S7. Retention of the first intron in *NRPA2* mRNA**

Analysis of intron retention in *NRPA2* as indicated in Figure 3 for *HD2B*.

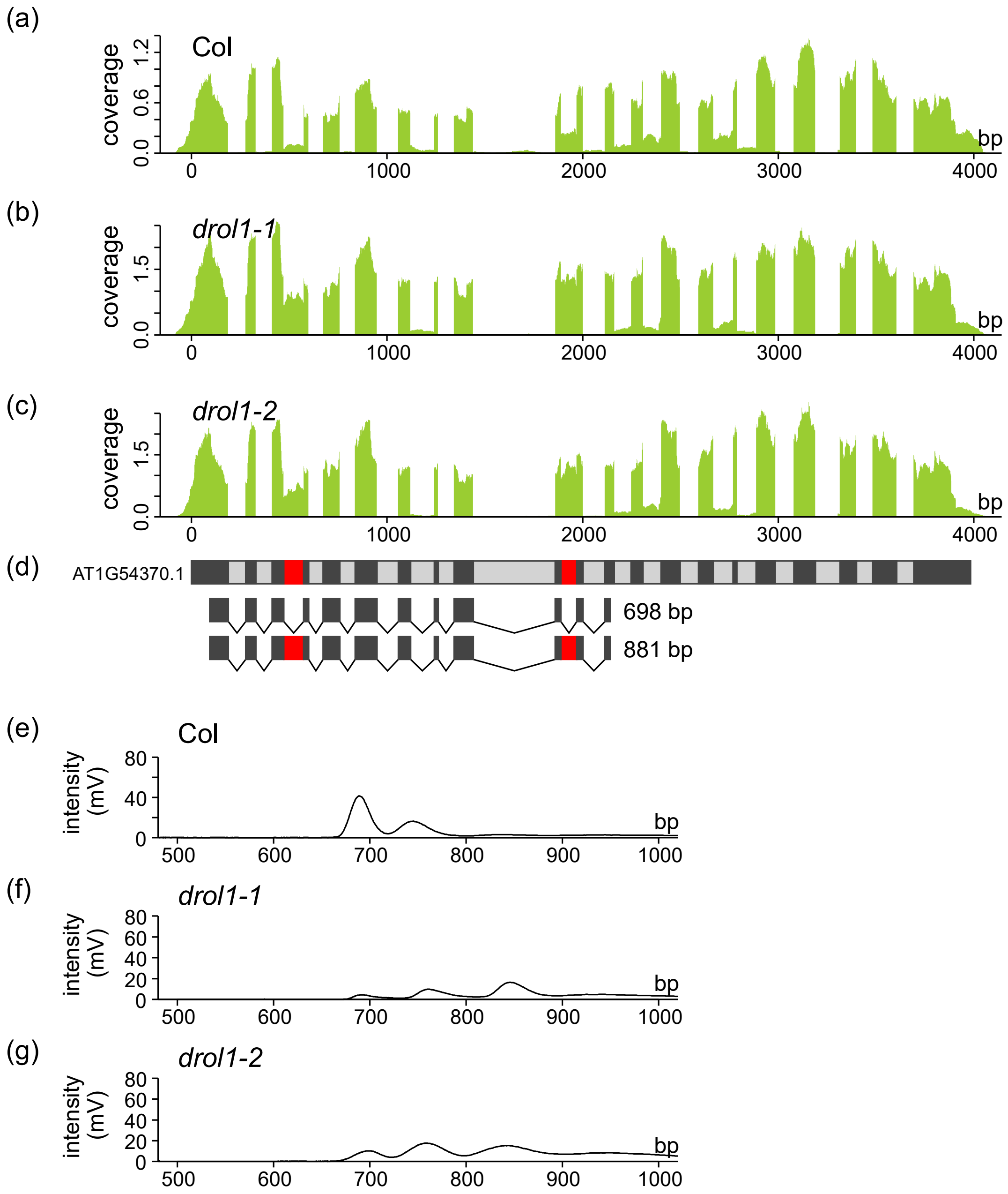


**Figure S8. Retention of the third and eleventh introns in *NHX5* mRNA**

Analysis of intron retention in *NHX5* as indicated in Figure 3 for *HD2B*.

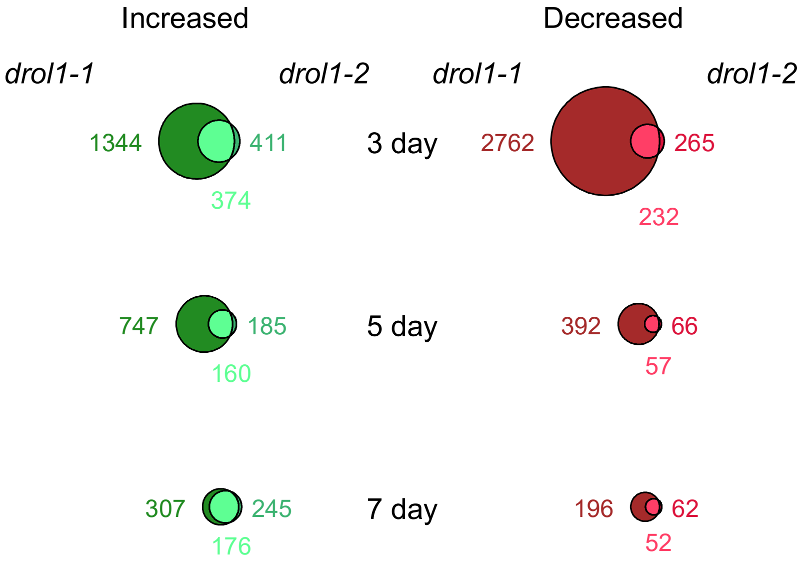


**Figure S9. Differentially expressed genes (DEGs) in *drol1* mutants**

DEGs were counted and displayed in circles. Green and red circles indicate upregulated and downregulated genes, respectively, in seedlings of *drol1* mutants when compared with the wild-type. Genes are also indicated by the numbers beside the circles and overlaps.

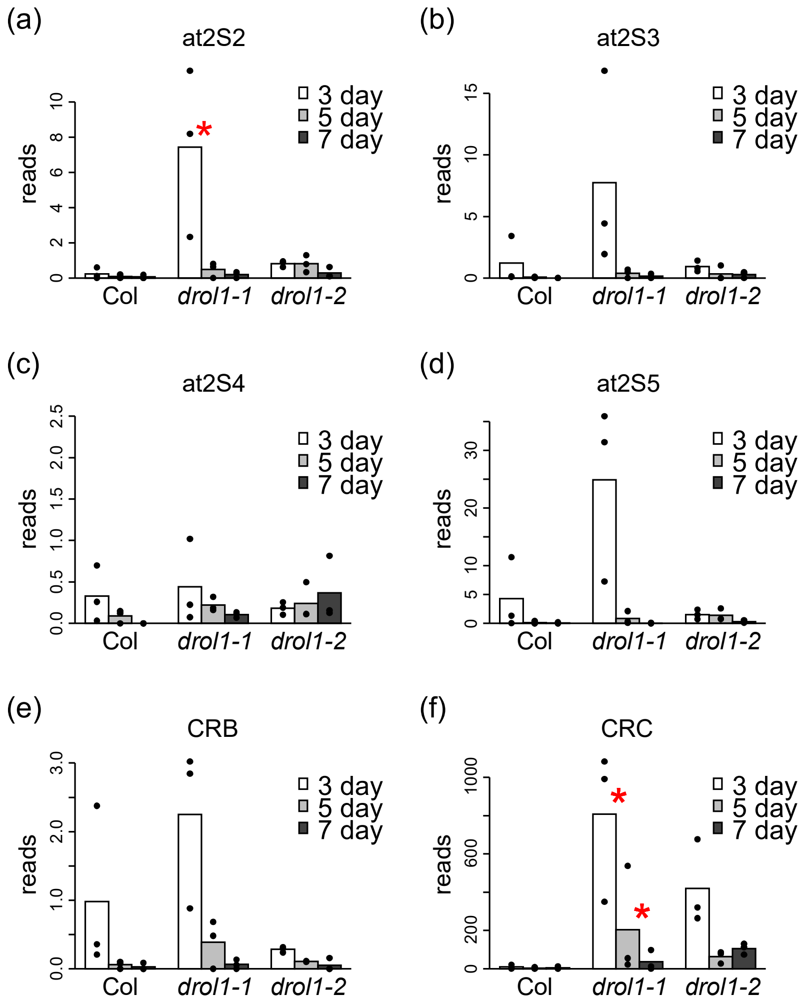


**Figure S10. Increased expression of genes encoding seed-storage proteins in *drol1* seedlings**

Expression levels of (a–d) four 2S albumin and (e–f) two 18S globulin genes determined by RNA-Seq. Charts and legends are the same as those shown in Figure 5.

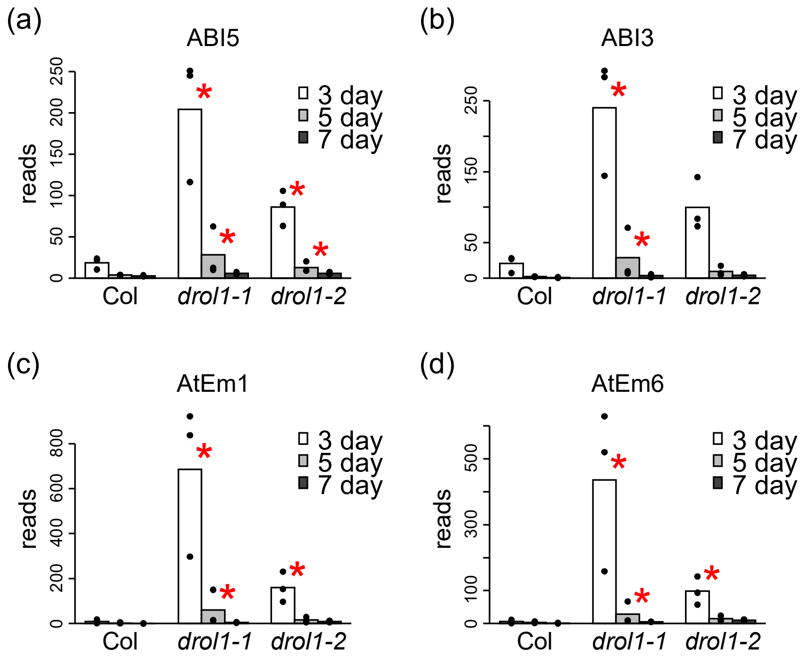


**Figure S11. Increased expression of genes responsible to ABA in *drol1* seedlings**

Expression levels of (a) *ABI5*, (b) *ABI3*, (c) *AtEm1*, and (d) *AtEm6* genes determined by RNA-Seq. Charts and legends are the same as those shown in Figure 5.

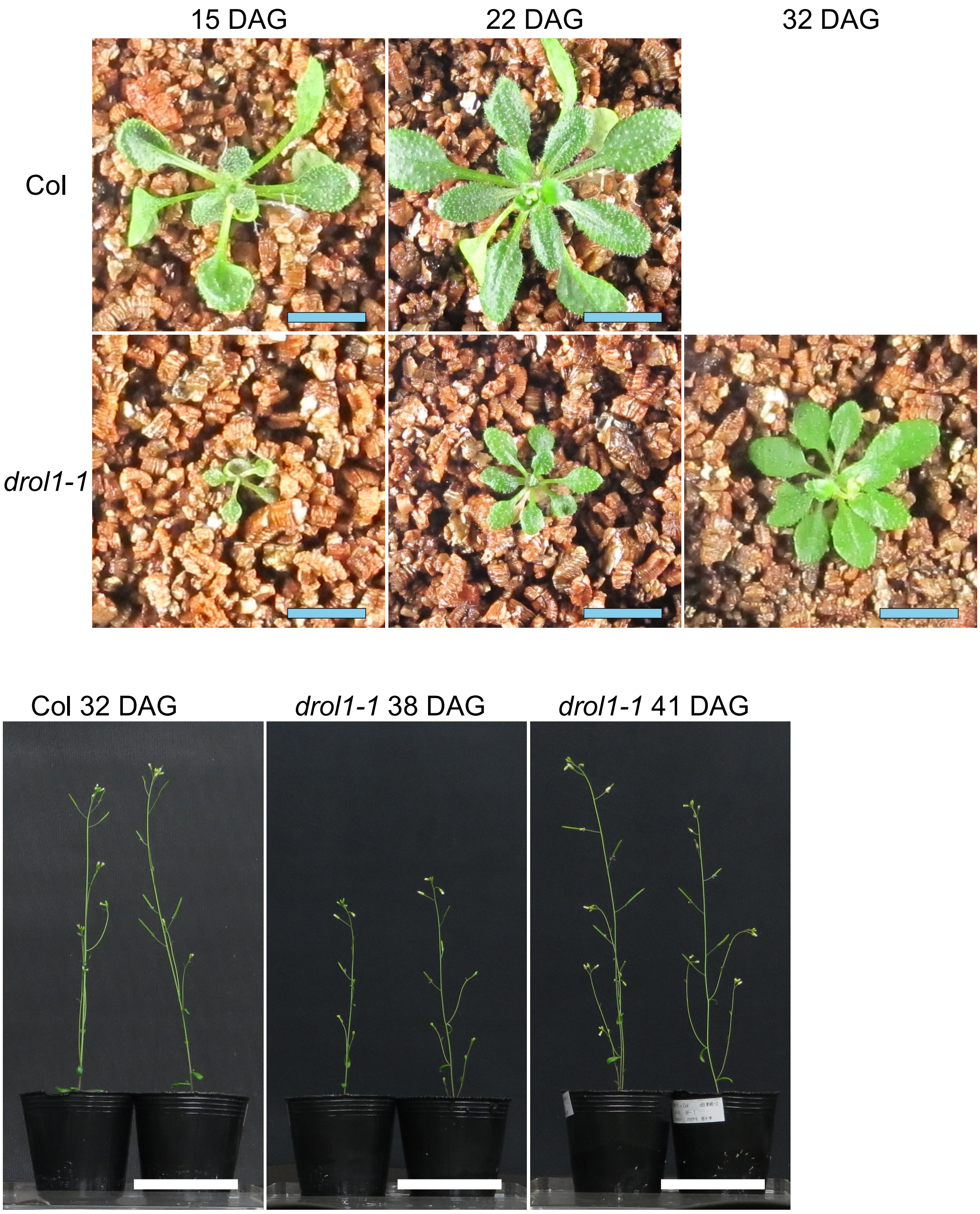


**Figure S12. Plant phenotypes of *drol1-1* mutants grown on soil**

After 2 weeks on growth medium, Col and *drol1-1* were planted in the soil. Photographs were taken on specific days after germination (DAG) as indicated. Blue and white bars indicate 1 cm and 5 cm, respectively.

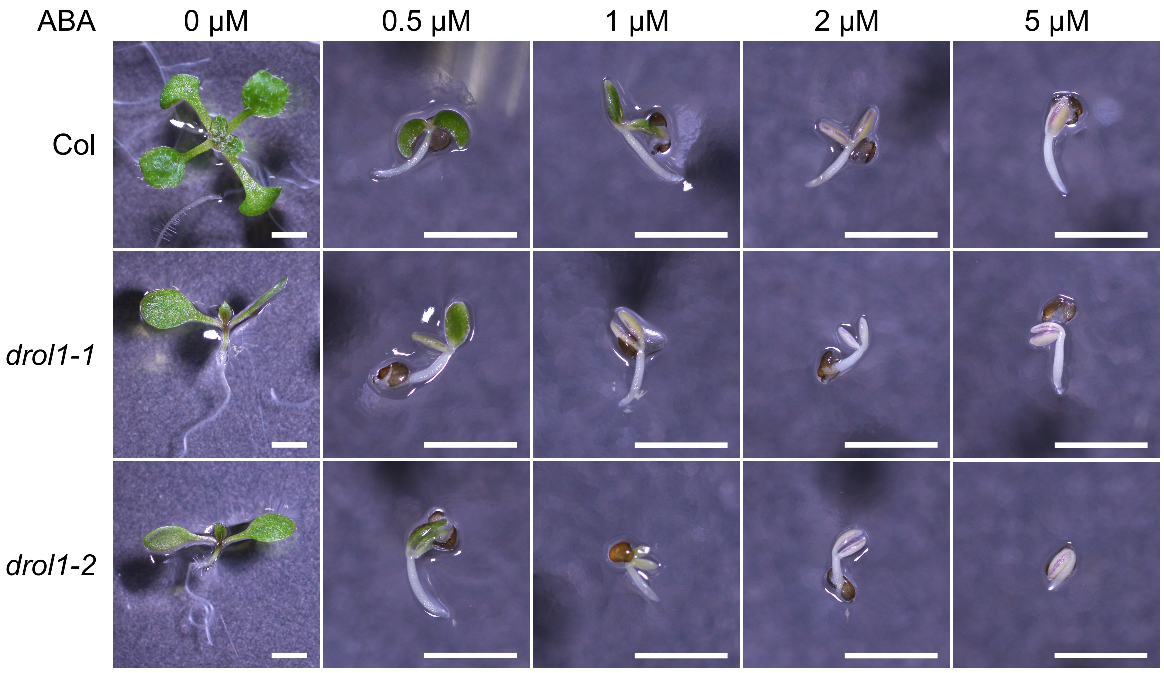


**Figure S13. Seedlings germinated on medium containing ABA**

Seeds for wild-type (Col), *drol1-1*, and *drol1-2* were placed on the medium containing ABA at various concentration as indicated. After 10 days of incubation in the growth chamber, seedlings were observed under microscopy. Bars = 1 mm.

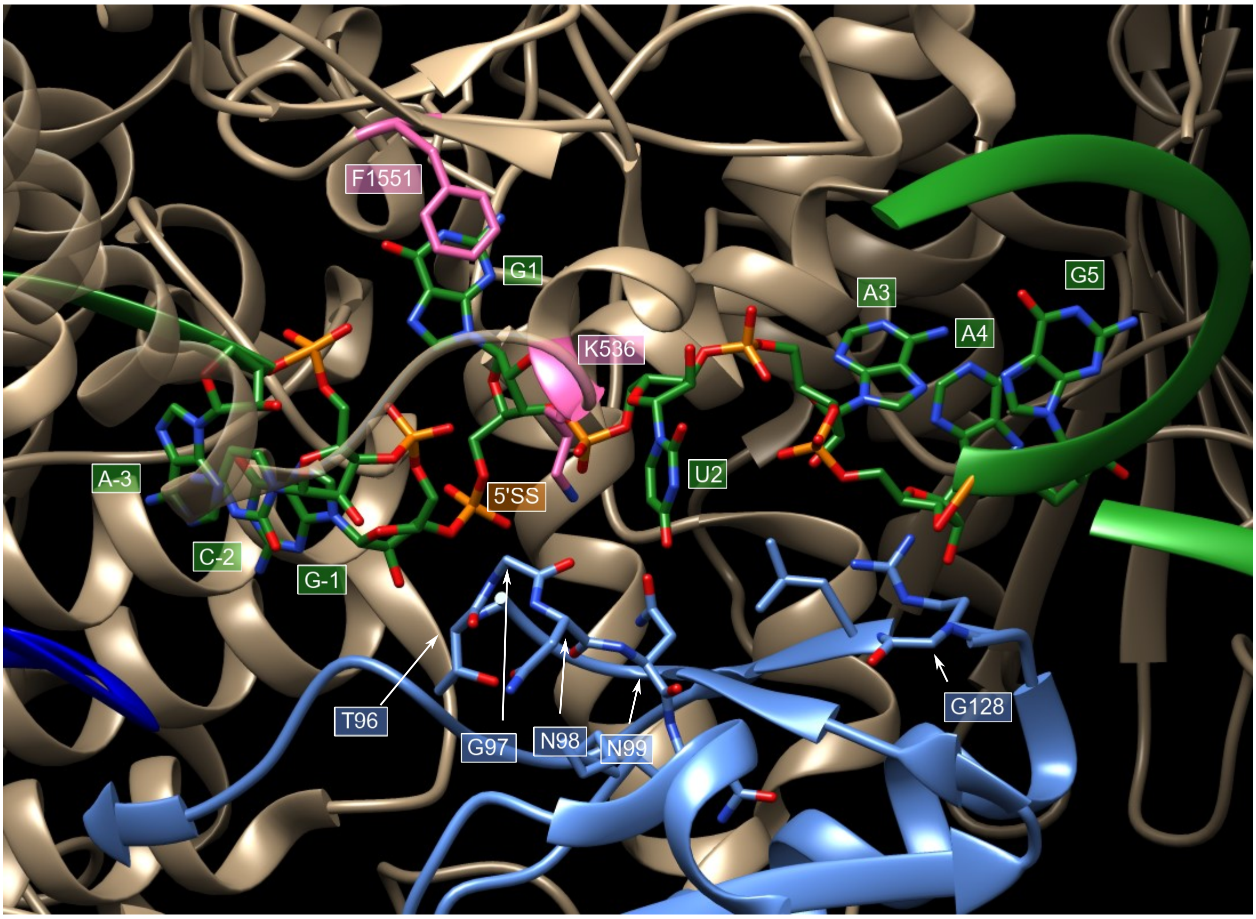


**Figure S14. Interaction between hDim1 and pre-mRNA**

The image was drawn based on original data (https://www.rcsb.org/structure/6AHD) (Zhan *et al.*, 2018) using UCSF Chimera (https://www.cgl.ucsf.edu/chimera/). hDim1, pre-mRNA, and PRP8 are represented in cyan, green, and beige, respectively.

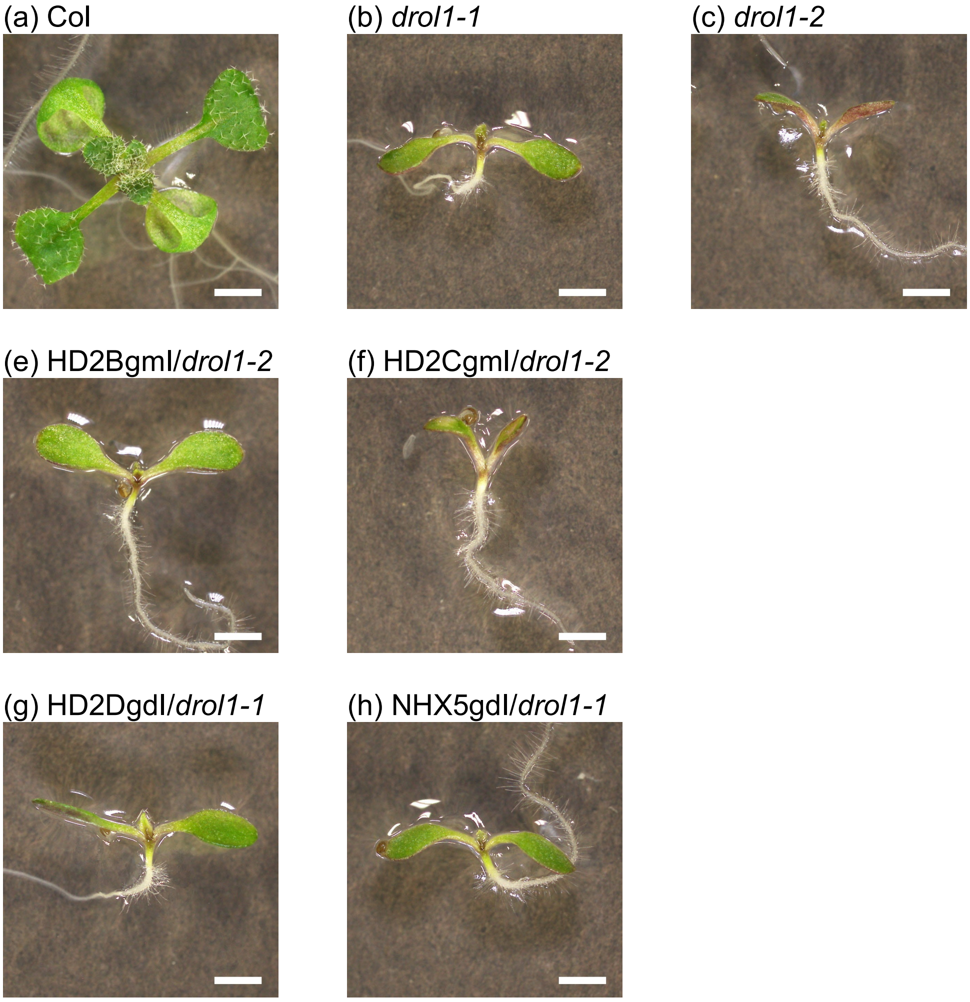


**Figure S15. Images for the drol1 mutants transformed with intron-modified genes**

(a-c) Images for 10-day-old seedlings for Col, *drol1-1*, and *drol1-2*, respectively. (d, e) *drol1-2* was transformed with the HD2B and HD2C in which AT–AC-type introns were substituted to GT–AG, respectively. (f, g) *drol1-1* was transformed with the HD2D and NHX5 gene lacking the AT–AC-type introns, respectively. Bars = 1mm.
