## Supplementary material for "*DROL1* subunit of U5 snRNP in the spliceosome is specifically required to splice AT–AC-type introns in *Arabidopsis*": Table S2

Table S2. Upregulated genes in *drol1* seedlings at 3 days after germination

| Gene ID | logFC | | Gene function | GO ID |
| --- | --- | --- | --- | --- |
|  | *drol1-1* | *drol1-2* |  |  |
| AT2G19320 | 9.23 | 6.34 | unknown protein |  |
| AT4G27140 | 9.03 | 7.26 | seed storage albumin 1 |  |
| AT4G31830 | 7.88 | 5.06 | unknown protein |  |
| AT2G28490 | 7.44 | 5.50 | RmlC-like cupins superfamily protein |  |
| AT3G22490 | 7.36 | 4.85 | Seed maturation protein |  |
| AT4G09600 | 7.20 | 5.26 | GAST1 protein homolog 3 |  |
| AT2G42560 | 7.15 | 5.39 | late embryogenesis abundant domain-containing protein / LEA domain-containing protein |  |
| AT5G44120 | 7.15 | 5.98 | RmlC-like cupins superfamily protein | GO:0009737 |
| AT5G62800 | 7.14 | 4.24 | Protein with RING/U-box and TRAF-like domains |  |
| AT3G15280 | 7.09 | 4.77 | unknown protein |  |
| AT3G24220 | 7.07 | 5.80 | nine-cis-epoxycarotenoid dioxygenase 6 |  |
| AT4G39130 | 7.05 | 4.99 | Dehydrin family protein |  |
| AT5G09640 | 7.04 | 5.36 | serine carboxypeptidase-like 19 |  |
| AT5G40420 | 7.00 | 5.01 | oleosin 2 | GO:0019915 |
| AT2G05420 | 6.91 | 6.71 | TRAF-like family protein |  |
| AT2G15010 | 6.91 | 5.70 | Plant thionin |  |
| AT5G10000 | 6.89 | 4.89 | ferredoxin 4 |  |
| AT1G52680 | 6.86 | 4.66 | late embryogenesis abundant protein-related / LEA protein-related |  |
| AT1G64295 | 6.79 | 5.23 | F-box associated ubiquitination effector family protein |  |
| AT3G27660 | 6.78 | 4.91 | oleosin 4 | GO:0019915 |
| AT1G27990 | 6.72 | 4.90 | unknown protein |  |
| AT2G44470 | 6.67 | 5.67 | beta glucosidase 29 | GO:0009651 |
| AT1G32710 | 6.64 | 4.67 | Cytochrome c oxidase, subunit Vib family protein |  |
| AT5G40382 | 6.62 | 5.13 | Cytochrome c oxidase subunit Vc family protein |  |
| AT5G01300 | 6.59 | 4.76 | PEBP (phosphatidylethanolamine-binding protein) family protein |  |
| AT2G34870 | 6.51 | 5.22 | hydroxyproline-rich glycoprotein family protein |  |
| AT4G25580 | 6.39 | 4.56 | CAP160 protein |  |
| AT3G29580 | 6.37 | 3.70 | Arabidopsis phospholipase-like protein (PEARLI 4) with TRAF-like domain |  |
| AT1G80090 | 6.36 | 4.69 | Cystathionine beta-synthase (CBS) family protein |  |
| AT1G04560 | 6.34 | 4.60 | AWPM-19-like family protein |  |
| AT1G07645 | 6.28 | 3.99 | dessication-induced 1VOC superfamily protein |  |
| AT5G22470 | 6.24 | 4.32 | NAD+ ADP-ribosyltransferases;NAD+ ADP-ribosyltransferases |  |
| AT2G18540 | 6.18 | 4.60 | RmlC-like cupins superfamily protein |  |
| AT3G51810 | 6.15 | 4.05 | Stress induced protein | GO:0009737 |
| AT1G48130 | 6.15 | 4.17 | 1-cysteine peroxiredoxin 1 | GO:0009414 |
| AT2G19900 | 6.15 | 4.56 | NADP-malic enzyme 1 |  |
| AT5G55240 | 6.11 | 4.18 | ARABIDOPSIS THALIANA PEROXYGENASE 2 |  |
| AT3G58450 | 6.11 | 3.95 | Adenine nucleotide alpha hydrolases-like superfamily protein |  |
| AT2G40170 | 6.11 | 3.96 | Stress induced protein | GO:0009737 |
| AT5G51760 | 6.09 | 3.86 | Protein phosphatase 2C family protein | GO:0009737 |
| AT2G21490 | 6.09 | 4.25 | dehydrin LEA | GO:0009414 |
|  |  |  |  | GO:0009737 |
| AT3G13280 | 6.04 | 6.18 | Putative endonuclease or glycosyl hydrolase |  |
| AT5G52300 | 6.03 | 4.52 | CAP160 protein | GO:0009414 |
|  |  |  |  | GO:0009737 |
|  |  |  |  | GO:0009651 |
| AT1G73190 | 6.02 | 4.15 | Aquaporin-like superfamily protein |  |
| AT4G36600 | 5.92 | 4.32 | Late embryogenesis abundant (LEA) protein |  |
| AT3G17520 | 5.87 | 4.38 | Late embryogenesis abundant protein (LEA) family protein |  |
| AT2G21820 | 5.82 | 3.90 | unknown protein |  |
| AT4G18920 | 5.81 | 3.83 | Protein of unknown function (DUF1264) |  |
| AT3G01570 | 5.80 | 3.85 | Oleosin family protein | GO:0019915 |
| AT5G07330 | 5.80 | 4.07 | unknown protein |  |
| AT2G23640 | 5.80 | 4.10 | Reticulan like protein B13 |  |
| AT1G48990 | 5.73 | 3.61 | Oleosin family protein |  |
| AT2G25890 | 5.72 | 3.96 | Oleosin family protein | GO:0019915 |
| AT1G02700 | 5.68 | 3.79 | unknown protein |  |
| AT1G52690 | 5.66 | 4.48 | Late embryogenesis abundant protein (LEA) family protein | GO:0009414 |
| AT3G03620 | 5.62 | 3.88 | MATE efflux family protein |  |
| AT2G47770 | 5.61 | 3.84 | TSPO(outer membrane tryptophan-rich sensory protein)-related | GO:0009737 |
|  |  |  |  | GO:0009651 |
| AT3G21370 | 5.60 | 4.47 | beta glucosidase 19 | GO:0009651 |
| AT3G54940 | 5.60 | 3.84 | Papain family cysteine protease |  |
| AT1G72100 | 5.57 | 3.84 | late embryogenesis abundant domain-containing protein / LEA domain-containing protein |  |
| AT5G45690 | 5.55 | 3.71 | Protein of unknown function (DUF1264) |  |
| AT3G03341 | 5.51 | 4.00 | unknown protein |  |
| AT3G53040 | 5.48 | 3.71 | late embryogenesis abundant protein, putative / LEA protein, putative |  |
| AT2G38905 | 5.47 | 3.55 | Low temperature and salt responsive protein family |  |
| AT5G65165 | 5.47 | 3.83 | succinate dehydrogenase 2-3 |  |
| AT3G15670 | 5.46 | 3.93 | Late embryogenesis abundant protein (LEA) family protein |  |
| AT5G16460 | 5.45 | 3.11 | Putative adipose-regulatory protein (Seipin) | GO:0019915 |
| AT2G36640 | 5.37 | 3.73 | embryonic cell protein 63 |  |
| AT1G56600 | 5.37 | 3.52 | galactinol synthase 2 | GO:0009414 |
|  |  |  |  | GO:0009737 |
|  |  |  |  | GO:0009651 |
| AT2G18340 | 5.37 | 3.83 | late embryogenesis abundant domain-containing protein / LEA domain-containing protein |  |
| AT5G04010 | 5.32 | 3.84 | F-box family protein |  |
| AT5G44310 | 5.29 | 4.10 | Late embryogenesis abundant protein (LEA) family protein |  |
| AT4G36700 | 5.29 | 4.19 | RmlC-like cupins superfamily protein |  |
| AT5G62490 | 5.28 | 3.32 | HVA22 homologue B | GO:0009737 |
|  |  |  |  | GO:0009651 |
| AT4G21020 | 5.25 | 4.09 | Late embryogenesis abundant protein (LEA) family protein |  |
| AT1G17810 | 5.24 | 3.49 | beta-tonoplast intrinsic protein |  |
| AT5G18450 | 5.24 | 3.87 | Integrase-type DNA-binding superfamily protein |  |
| AT5G43770 | 5.20 | 3.08 | proline-rich family protein |  |
| AT2G27940 | 5.17 | 4.04 | RING/U-box superfamily protein |  |
| AT4G16820 | 5.16 | 4.24 | alpha/beta-Hydrolases superfamily protein |  |
| AT5G42290 | 5.07 | 3.32 | transcription activator-related |  |
| AT2G29380 | 5.04 | 3.09 | highly ABA-induced PP2C gene 3 |  |
| AT5G49120 | 5.04 | 3.52 | Protein of unknown function (DUF581) |  |
| AT1G23070 | 5.03 | 3.62 | Protein of unknown function (DUF300) |  |
| AT1G03120 | 4.99 | 3.27 | responsive to abscisic acid 28 |  |
| AT1G16850 | 4.99 | 3.43 | unknown protein |  |
| AT4G27530 | 4.92 | 3.45 | unknown protein |  |
| AT5G47130 | 4.91 | 4.15 | Bax inhibitor-1 family protein |  |
| AT5G64210 | 4.89 | 3.19 | alternative oxidase 2 |  |
| AT1G14930 | 4.87 | 3.17 | Polyketide cyclase/dehydrase and lipid transport superfamily protein |  |
| AT1G01520 | 4.86 | 3.86 | Homeodomain-like superfamily protein |  |
| AT1G04660 | 4.85 | 3.79 | glycine-rich protein |  |
| AT5G06760 | 4.80 | 3.43 | Late Embryogenesis Abundant 4-5 | GO:0009414 |
| AT1G03790 | 4.78 | 3.08 | Zinc finger C-x8-C-x5-C-x3-H type family protein |  |
| AT3G18610 | 4.75 | 3.66 | nucleolin like 2 |  |
| AT5G05220 | 4.74 | 4.07 | unknown protein |  |
| AT4G18620 | 4.73 | 3.02 | PYR1-like 13 | GO:0009737 |
| AT1G32560 | 4.72 | 3.10 | Late embryogenesis abundant protein, group 1 protein | GO:0009414 |
| AT1G22600 | 4.69 | 3.08 | Late embryogenesis abundant protein (LEA) family protein |  |
| AT5G44260 | 4.64 | 2.56 | Zinc finger C-x8-C-x5-C-x3-H type family protein |  |
| AT1G67856 | 4.61 | 3.83 | RING/U-box superfamily protein |  |
| AT3G50980 | 4.60 | 3.16 | dehydrin xero 1 | GO:0009414 |
|  |  |  |  | GO:0009737 |
| AT3G11050 | 4.60 | 3.07 | ferritin 2 | GO:0009737 |
| AT2G18050 | 4.58 | 3.69 | histone H1-3 |  |
| AT2G02850 | 4.58 | 2.88 | plantacyanin |  |
| AT4G01970 | 4.54 | 2.85 | stachyose synthase |  |
| AT1G47980 | 4.54 | 3.69 | unknown protein |  |
| AT5G51210 | 4.54 | 2.84 | oleosin3 | GO:0019915 |
| AT1G24580 | 4.52 | 3.25 | RING/U-box superfamily protein |  |
| AT1G67855 | 4.51 | 3.80 | unknown protein |  |
| AT5G24130 | 4.51 | 2.92 | unknown protein |  |
| AT2G31980 | 4.50 | 2.76 | PHYTOCYSTATIN 2 |  |
| AT5G01670 | 4.50 | 2.57 | NAD(P)-linked oxidoreductase superfamily protein |  |
| AT1G16730 | 4.50 | 2.85 | unknown protein 6 |  |
| AT4G11910 | 4.47 | 3.30 | unknown protein |  |
| AT5G63350 | 4.47 | 3.19 | unknown protein |  |
| AT1G04920 | 4.44 | 2.95 | sucrose phosphate synthase 3F |  |
| AT4G10020 | 4.44 | 3.21 | hydroxysteroid dehydrogenase 5 |  |
| AT3G23860 | 4.43 | 3.83 | GTP-binding protein-related |  |
| AT5G59390 | 4.38 | 3.95 | XH/XS domain-containing protein |  |
| AT1G15330 | 4.30 | 3.28 | Cystathionine beta-synthase (CBS) protein |  |
| AT2G33280 | 4.30 | 3.75 | Major facilitator superfamily protein |  |
| AT2G21720 | 4.28 | 2.75 | Plant protein of unknown function (DUF639) |  |
| AT5G13210 | 4.28 | 3.19 | Uncharacterised conserved protein UCP015417, vWA |  |
| AT2G32830 | 4.25 | 2.64 | phosphate transporter 1;5 |  |
| AT4G15396 | 4.22 | 4.09 | cytochrome P450, family 702, subfamily A, polypeptide 6 |  |
| AT1G67100 | 4.19 | 3.04 | LOB domain-containing protein 40 |  |
| AT3G25260 | 4.14 | 2.80 | Major facilitator superfamily protein |  |
| AT1G80740 | 4.05 | 3.52 | chromomethylase 1 |  |
| AT5G57790 | 4.05 | 2.86 | unknown protein |  |
| AT4G02690 | 4.05 | 2.51 | Bax inhibitor-1 family protein |  |
| AT5G60760 | 4.04 | 2.65 | P-loop containing nucleoside triphosphate hydrolases superfamily protein |  |
| AT4G24000 | 4.03 | 4.26 | cellulose synthase like G2 |  |
| AT3G48510 | 4.02 | 3.19 | unknown protein |  |
| AT5G22545 | 4.02 | 3.49 | unknown protein |  |
| AT5G01520 | 4.01 | 2.81 | RING/U-box superfamily protein | GO:0009737 |
|  |  |  |  | GO:0009651 |
| AT2G35590 | 3.98 | 4.40 | pseudogene |  |
| AT4G33467 | 3.94 | 3.56 | unknown protein |  |
| AT4G15380 | 3.94 | 2.31 | cytochrome P450, family 705, subfamily A, polypeptide 4 |  |
| AT2G18180 | 3.93 | 3.88 | Sec14p-like phosphatidylinositol transfer family protein |  |
| AT3G20710 | 3.92 | 2.47 | F-box family protein |  |
| AT5G53870 | 3.90 | 2.84 | early nodulin-like protein 1 |  |
| AT2G18570 | 3.86 | 2.55 | UDP-Glycosyltransferase superfamily protein |  |
| AT5G04500 | 3.86 | 2.41 | glycosyltransferase family protein 47 | GO:0009737 |
|  |  |  |  | GO:0009651 |
| AT3G21380 | 3.86 | 3.60 | Mannose-binding lectin superfamily protein |  |
| AT1G22340 | 3.81 | 2.88 | UDP-glucosyl transferase 85A7 |  |
| AT1G35910 | 3.80 | 2.83 | Haloacid dehalogenase-like hydrolase (HAD) superfamily protein | GO:0009651 |
| AT3G51750 | 3.78 | 2.28 | unknown protein |  |
| AT4G18650 | 3.78 | 2.55 | transcription factor-related |  |
| AT5G59320 | 3.77 | 2.37 | lipid transfer protein 3 | GO:0009414 |
|  |  |  |  | GO:0009737 |
| AT4G02280 | 3.76 | 2.24 | sucrose synthase 3 | GO:0009414 |
| AT2G17680 | 3.71 | 2.79 | Arabidopsis protein of unknown function (DUF241) |  |
| AT5G50360 | 3.69 | 2.89 | unknown protein |  |
| AT2G38465 | 3.68 | 2.27 | unknown protein |  |
| AT2G35570 | 3.67 | 4.00 | pseudogene |  |
| AT5G38780 | 3.67 | 2.63 | S-adenosyl-L-methionine-dependent methyltransferases superfamily protein |  |
| AT2G15130 | 3.66 | 3.47 | Plant basic secretory protein (BSP) family protein |  |
| AT5G57550 | 3.64 | 3.18 | xyloglucan endotransglucosylase/hydrolase 25 |  |
| AT2G27380 | 3.64 | 3.78 | extensin proline-rich 1 |  |
| AT1G64110 | 3.63 | 2.74 | P-loop containing nucleoside triphosphate hydrolases superfamily protein |  |
| AT3G24542 | 3.62 | 3.80 | Beta-galactosidase related protein |  |
| AT1G21520 | 3.62 | 2.47 | unknown protein |  |
| AT5G50770 | 3.60 | 2.74 | hydroxysteroid dehydrogenase 6 |  |
| AT1G07430 | 3.60 | 2.05 | highly ABA-induced PP2C gene 2 |  |
| AT4G12680 | 3.59 | 2.40 | unknown protein |  |
| AT3G44830 | 3.58 | 2.14 | Lecithin:cholesterol acyltransferase family protein |  |
| AT1G77120 | 3.57 | 2.36 | alcohol dehydrogenase 1 | GO:0009414 |
|  |  |  |  | GO:0009737 |
|  |  |  |  | GO:0009651 |
| AT4G15390 | 3.57 | 2.20 | HXXXD-type acyl-transferase family protein |  |
| AT4G01180 | 3.56 | 3.42 | XH/XS domain-containing protein |  |
| AT2G18190 | 3.56 | 3.00 | P-loop containing nucleoside triphosphate hydrolases superfamily protein | GO:0009651 |
| AT2G04050 | 3.55 | 3.07 | MATE efflux family protein |  |
| AT5G03795 | 3.50 | 2.12 | Exostosin family protein |  |
| AT5G55470 | 3.49 | 2.26 | Na+/H+ (sodium hydrogen) exchanger 3 |  |
| AT2G36270 | 3.47 | 2.22 | Basic-leucine zipper (bZIP) transcription factor family protein | GO:0009414 |
|  |  |  |  | GO:0009737 |
|  |  |  |  | GO:0009651 |
| AT3G22860 | 3.43 | 3.74 | eukaryotic translation initiation factor 3 subunit C2 |  |
| AT5G55460 | 3.39 | 2.17 | Bifunctional inhibitor/lipid-transfer protein/seed storage 2S albumin superfamily protein |  |
| AT4G19810 | 3.37 | 2.61 | Glycosyl hydrolase family protein with chitinase insertion domain | GO:0009737 |
|  |  |  |  | GO:0009651 |
| AT1G78780 | 3.36 | 2.01 | pathogenesis-related family protein |  |
| AT2G05915 | 3.33 | 3.01 | unknown protein |  |
| AT3G14880 | 3.30 | 2.26 | unknown protein |  |
| AT2G20800 | 3.29 | 1.91 | NAD(P)H dehydrogenase B4 |  |
| AT3G27473 | 3.27 | 2.10 | Cysteine/Histidine-rich C1 domain family protein |  |
| AT5G45310 | 3.24 | 2.12 | unknown protein |  |
| AT1G01240 | 3.18 | 1.96 | unknown protein |  |
| AT4G18220 | 3.16 | 2.53 | Drug/metabolite transporter superfamily protein |  |
| AT5G18220 | 3.11 | 2.43 | O-Glycosyl hydrolases family 17 protein |  |
| AT2G43960 | 3.11 | 3.01 | SWAP (Suppressor-of-White-APricot)/surp domain-containing protein |  |
| AT1G65370 | 3.09 | 2.73 | TRAF-like family protein |  |
| AT2G02120 | 3.06 | 1.78 | Scorpion toxin-like knottin superfamily protein |  |
| AT3G30460 | 3.05 | 2.33 | RING/U-box superfamily protein |  |
| AT4G24040 | 3.04 | 2.19 | trehalase 1 |  |
| AT5G28235 | 3.03 | 3.13 | Ulp1 protease family protein |  |
| AT2G04070 | 3.03 | 2.54 | MATE efflux family protein |  |
| AT3G45730 | 3.03 | 2.06 | unknown protein |  |
| AT4G12290 | 3.02 | 1.45 | Copper amine oxidase family protein |  |
| AT5G52310 | 2.99 | 2.21 | low-temperature-responsive protein 78 (LTI78) / desiccation-responsive protein 29A (RD29A) | GO:0009414 |
|  |  |  |  | GO:0009737 |
|  |  |  |  | GO:0009651 |
| AT4G23600 | 2.99 | 3.75 | Tyrosine transaminase family protein | GO:0009737 |
|  |  |  |  | GO:0009651 |
| AT3G06435 | 2.98 | 2.41 | Expressed protein |  |
| AT5G59340 | 2.98 | 2.28 | WUSCHEL related homeobox 2 |  |
| AT1G19490 | 2.98 | 1.90 | Basic-leucine zipper (bZIP) transcription factor family protein |  |
| AT5G45570 | 2.96 | 3.01 | Ulp1 protease family protein |  |
| AT4G35690 | 2.96 | 3.06 | Arabidopsis protein of unknown function (DUF241) |  |
| AT1G08830 | 2.95 | 2.49 | copper/zinc superoxide dismutase 1 | GO:0009651 |
| AT1G62710 | 2.94 | 1.85 | beta vacuolar processing enzyme |  |
| AT3G28500 | 2.94 | 2.33 | 60S acidic ribosomal protein family |  |
| AT5G07060 | 2.92 | 2.38 | CCCH-type zinc fingerfamily protein with RNA-binding domain |  |
| AT1G62290 | 2.92 | 1.85 | Saposin-like aspartyl protease family protein |  |
| AT3G30120 | 2.91 | 2.27 | pseudogene |  |
| AT1G07985 | 2.87 | 2.26 | Expressed protein |  |
| AT3G12203 | 2.85 | 2.04 | serine carboxypeptidase-like 17 |  |
| AT5G58610 | 2.82 | 2.47 | PHD finger transcription factor, putative |  |
| AT4G10440 | 2.82 | 3.29 | S-adenosyl-L-methionine-dependent methyltransferases superfamily protein |  |
| AT5G22110 | 2.81 | 2.56 | DNA polymerase epsilon subunit B2 |  |
| AT3G21600 | 2.80 | 1.26 | Senescence/dehydration-associated protein-related |  |
| AT2G19850 | 2.80 | 3.35 | unknown protein |  |
| AT5G44417 | 2.79 | 2.33 | pseudogene |  |
| AT2G28190 | 2.78 | 1.33 | copper/zinc superoxide dismutase 2 | GO:0009651 |
| AT2G34850 | 2.76 | 2.13 | NAD(P)-binding Rossmann-fold superfamily protein |  |
| AT5G06755 | 2.75 | 2.94 | unknown protein |  |
| AT3G24780 | 2.72 | 2.12 | Uncharacterised conserved protein UCP015417, vWA |  |
| AT1G14520 | 2.72 | 1.97 | myo-inositol oxygenase 1 |  |
| AT1G12520 | 2.70 | 1.51 | copper chaperone for SOD1 |  |
| AT5G47810 | 2.69 | 1.27 | phosphofructokinase 2 |  |
| AT5G39850 | 2.69 | 1.60 | Ribosomal protein S4 |  |
| AT3G15357 | 2.68 | 1.59 | unknown protein |  |
| AT1G01470 | 2.66 | 1.61 | Late embryogenesis abundant protein | GO:0009414 |
| AT2G43660 | 2.65 | 1.84 | Carbohydrate-binding X8 domain superfamily protein |  |
| AT5G02580 | 2.65 | 2.10 | Plant protein 1589 of unknown function |  |
| AT1G79610 | 2.65 | 2.02 | Na+/H+ antiporter 6 |  |
| AT4G36620 | 2.64 | 1.45 | GATA transcription factor 19 |  |
| AT3G01345 | 2.63 | 2.91 | Expressed protein |  |
| AT4G36105 | 2.63 | 2.61 | unknown protein |  |
| AT3G30720 | 2.59 | 2.90 | qua-quine starch |  |
| AT1G10300 | 2.58 | 1.59 | Nucleolar GTP-binding protein |  |
| AT1G69260 | 2.55 | 1.76 | ABI five binding protein | GO:0009737 |
| AT2G15030 | 2.55 | 1.52 | pseudogene |  |
| AT5G06980 | 2.53 | 2.19 | unknown protein |  |
| AT3G01600 | 2.52 | 1.64 | NAC domain containing protein 44 |  |
| AT4G31520 | 2.52 | 1.89 | SDA1 family protein |  |
| AT4G26590 | 2.48 | 1.55 | oligopeptide transporter 5 |  |
| AT3G28890 | 2.47 | 1.58 | receptor like protein 43 |  |
| AT1G26800 | 2.46 | 1.64 | RING/U-box superfamily protein |  |
| AT4G22920 | 2.45 | 1.80 | non-yellowing 1 |  |
| AT4G23050 | 2.44 | 1.34 | PAS domain-containing protein tyrosine kinase family protein |  |
| AT5G22290 | 2.44 | 1.44 | NAC domain containing protein 89 |  |
| AT5G07010 | 2.43 | 2.10 | sulfotransferase 2A |  |
| AT5G45095 | 2.43 | 1.53 | unknown protein |  |
| AT1G53480 | 2.41 | 1.98 | mto 1 responding down 1 |  |
| AT4G04030 | 2.39 | 3.00 | ovate family protein 9 |  |
| AT5G66580 | 2.37 | 1.45 | unknown protein |  |
| AT2G39820 | 2.32 | 3.09 | Translation initiation factor IF6 |  |
| AT5G65890 | 2.31 | 1.31 | ACT domain repeat 1 |  |
| AT1G13740 | 2.29 | 1.43 | ABI five binding protein 2 | GO:0009414 |
|  |  |  |  | GO:0009737 |
| AT4G24480 | 2.28 | 1.48 | Protein kinase superfamily protein |  |
| AT2G43590 | 2.28 | 1.78 | Chitinase family protein |  |
| AT1G29940 | 2.27 | 2.21 | nuclear RNA polymerase A2 |  |
| AT5G53700 | 2.27 | 2.08 | RNA-binding (RRM/RBD/RNP motifs) family protein |  |
| AT5G44980 | 2.26 | 1.74 | F-box/RNI-like/FBD-like domains-containing protein |  |
| AT4G22990 | 2.25 | 1.56 | Major Facilitator Superfamily with SPX (SYG1/Pho81/XPR1) domain-containing protein |  |
| AT3G22770 | 2.25 | 2.88 | F-box associated ubiquitination effector family protein |  |
| AT5G60142 | 2.25 | 2.81 | AP2/B3-like transcriptional factor family protein |  |
| AT3G61630 | 2.25 | 1.70 | cytokinin response factor 6 |  |
| AT4G22470 | 2.23 | 2.50 | protease inhibitor/seed storage/lipid transfer protein (LTP) family protein |  |
| AT3G54730 | 2.23 | 2.82 | unknown protein |  |
| AT2G34655 | 2.22 | 2.16 | unknown protein |  |
| AT3G13090 | 2.20 | 2.07 | multidrug resistance-associated protein 8 |  |
| AT5G45540 | 2.18 | 1.76 | Protein of unknown function (DUF594) |  |
| AT4G00390 | 2.18 | 1.99 | DNA-binding storekeeper protein-related transcriptional regulator |  |
| AT2G18193 | 2.18 | 1.57 | P-loop containing nucleoside triphosphate hydrolases superfamily protein |  |
| AT5G56100 | 2.17 | 1.88 | glycine-rich protein / oleosin |  |
| AT5G02200 | 2.15 | 2.30 | far-red-elongated hypocotyl1-like |  |
| AT3G12320 | 2.14 | 1.58 | unknown protein |  |
| AT1G07180 | 2.13 | 1.67 | alternative NAD(P)H dehydrogenase 1 |  |
| AT1G08430 | 2.12 | 2.40 | aluminum-activated malate transporter 1 |  |
| AT5G61740 | 2.11 | 2.24 | ABC2 homolog 14 |  |
| AT1G71280 | 2.10 | 1.45 | DEA(D/H)-box RNA helicase family protein |  |
| AT4G25170 | 2.09 | 1.66 | Uncharacterised conserved protein (UCP012943) |  |
| AT5G18210 | 2.07 | 1.53 | NAD(P)-binding Rossmann-fold superfamily protein |  |
| AT4G01460 | 2.06 | 1.38 | basic helix-loop-helix (bHLH) DNA-binding superfamily protein |  |
| AT5G59670 | 2.05 | 1.85 | Leucine-rich repeat protein kinase family protein |  |
| AT2G46840 | 2.04 | 1.93 | DOMAIN OF UNKNOWN FUNCTION 724 4 |  |
| AT2G43570 | 2.04 | 1.48 | chitinase, putative |  |
| AT3G03310 | 2.04 | 1.00 | lecithin:cholesterol acyltransferase 3 |  |
| AT5G17800 | 2.02 | 1.37 | myb domain protein 56 |  |
| AT1G62530 | 2.02 | 1.79 | Plant protein of unknown function (DUF863) |  |
| AT2G18720 | 2.01 | 2.16 | Translation elongation factor EF1A/initiation factor IF2gamma family protein |  |
| AT5G61270 | 1.97 | 1.54 | phytochrome-interacting factor7 |  |
| AT1G12570 | 1.93 | 2.17 | Glucose-methanol-choline (GMC) oxidoreductase family protein |  |
| AT3G53210 | 1.93 | 1.48 | nodulin MtN21 /EamA-like transporter family protein |  |
| AT1G48400 | 1.92 | 2.23 | F-box/RNI-like/FBD-like domains-containing protein |  |
| AT1G08910 | 1.92 | 1.76 | zinc ion binding;zinc ion binding | GO:0009737 |
|  |  |  |  | GO:0009651 |
| AT3G27220 | 1.92 | 1.74 | Galactose oxidase/kelch repeat superfamily protein |  |
| AT3G05640 | 1.89 | 1.71 | Protein phosphatase 2C family protein | GO:0009414 |
| AT2G40010 | 1.88 | 1.26 | Ribosomal protein L10 family protein |  |
| AT5G24280 | 1.85 | 1.23 | gamma-irradiation and mitomycin c induced 1 |  |
| AT2G43500 | 1.85 | 1.22 | Plant regulator RWP-RK family protein |  |
| AT5G58670 | 1.84 | 1.59 | phospholipase C1 | GO:0009414 |
|  |  |  |  | GO:0009737 |
|  |  |  |  | GO:0009651 |
| AT3G62090 | 1.81 | 1.23 | phytochrome interacting factor 3-like 2 |  |
| AT5G60250 | 1.81 | 1.24 | zinc finger (C3HC4-type RING finger) family protein |  |
| AT3G05650 | 1.80 | 2.32 | receptor like protein 32 |  |
| AT4G00893 | 1.78 | 1.62 | unknown protein |  |
| AT5G66080 | 1.78 | 1.37 | Protein phosphatase 2C family protein |  |
| AT3G26165 | 1.75 | 1.90 | CYP71B18 (\"cytochrome P450, family 71, subfamily B, polypeptide 18\") |  |
| AT5G38200 | 1.74 | 1.46 | Class I glutamine amidotransferase-like superfamily protein |  |
| AT3G15790 | 1.74 | 1.08 | methyl-CPG-binding domain 11 |  |
| AT4G34790 | 1.74 | 1.61 | SAUR-like auxin-responsive protein family |  |
| AT5G08600 | 1.73 | 1.79 | U3 ribonucleoprotein (Utp) family protein |  |
| AT2G35950 | 1.72 | 2.61 | embryo sac development arrest 12 |  |
| AT1G67960 | 1.71 | 1.70 | unknown protein |  |
| AT5G41765 | 1.70 | 1.81 | DNA-binding storekeeper protein-related transcriptional regulator |  |
| AT1G15790 | 1.70 | 1.62 | unknown protein |  |
| AT3G28580 | 1.70 | 2.38 | P-loop containing nucleoside triphosphate hydrolases superfamily protein | GO:0009737 |
| AT4G39590 | 1.69 | 1.85 | Galactose oxidase/kelch repeat superfamily protein |  |
| AT1G76170 | 1.68 | 1.35 | 2-thiocytidine tRNA biosynthesis protein, TtcA |  |
| AT1G17960 | 1.68 | 1.43 | Threonyl-tRNA synthetase |  |
| AT4G31877 | 1.67 | 1.59 | MIR156C; miRNA |  |
| AT2G46270 | 1.65 | 1.36 | G-box binding factor 3 | GO:0009737 |
| AT1G30040 | 1.64 | 1.68 | gibberellin 2-oxidase |  |
| AT5G57240 | 1.64 | 1.14 | OSBP(oxysterol binding protein)-related protein 4C |  |
| AT1G35530 | 1.63 | 1.59 | DEAD/DEAH box RNA helicase family protein |  |
| AT1G34180 | 1.62 | 1.23 | NAC domain containing protein 16 |  |
| AT1G18100 | 1.62 | 1.22 | PEBP (phosphatidylethanolamine-binding protein) family protein | GO:0009737 |
| AT1G68945 | 1.59 | 1.36 | unknown protein |  |
| AT1G06720 | 1.59 | 1.18 | P-loop containing nucleoside triphosphate hydrolases superfamily protein |  |
| AT5G56900 | 1.58 | 1.31 | CwfJ-like family protein / zinc finger (CCCH-type) family protein |  |
| AT1G68870 | 1.58 | 1.05 | SOB five-like 2 |  |
| AT5G33390 | 1.52 | 1.20 | glycine-rich protein |  |
| AT4G24900 | 1.52 | 1.40 | unknown protein |  |
| AT1G65200 | 1.50 | 2.04 | Ubiquitin carboxyl-terminal hydrolase-related protein |  |
| AT5G25460 | 1.50 | 1.07 | Protein of unknown function, DUF642 |  |
| AT5G15960 | 1.50 | 1.76 | stress-responsive protein (KIN1) / stress-induced protein (KIN1) | GO:0009414 |
|  |  |  |  | GO:0009737 |
| AT3G48260 | 1.49 | 1.25 | with no lysine (K) kinase 3 |  |
| AT4G04745 | 1.45 | 1.14 | unknown protein |  |
| AT3G27620 | 1.43 | 1.21 | alternative oxidase 1C |  |
| AT3G06820 | 1.43 | 1.23 | Mov34/MPN/PAD-1 family protein |  |
| AT4G19960 | 1.41 | 1.29 | K+ uptake permease 9 |  |
| AT5G52570 | 1.40 | 1.04 | beta-carotene hydroxylase 2 |  |
| AT4G21326 | 1.38 | 1.39 | subtilase 3.12 |  |
| AT5G49300 | 1.33 | 1.04 | GATA transcription factor 16 |  |
| AT4G21680 | 1.32 | 1.37 | NITRATE TRANSPORTER 1.8 |  |
| AT3G53800 | 1.32 | 0.94 | Fes1B |  |
| AT1G32583 | 1.31 | 1.15 | unknown protein |  |
| AT1G66760 | 1.31 | 1.34 | MATE efflux family protein |  |
| AT5G65640 | 1.30 | 1.09 | beta HLH protein 93 |  |
| AT4G38810 | 1.30 | 0.81 | Calcium-binding EF-hand family protein |  |
| AT2G42660 | 1.25 | 1.52 | Homeodomain-like superfamily protein |  |
| AT4G15260 | 1.23 | 1.00 | UDP-Glycosyltransferase superfamily protein |  |
| AT2G37740 | 1.23 | 1.58 | zinc-finger protein 10 |  |
| AT5G57640 | 1.22 | 1.62 | GCK domain-containing protein |  |
| AT5G59660 | 1.22 | 1.06 | Leucine-rich repeat protein kinase family protein |  |
| AT1G77320 | 1.21 | 1.20 | transcription coactivators |  |
| AT3G59330 | 1.20 | 1.04 | Eukaryotic protein of unknown function (DUF914) |  |
| AT5G65610 | 1.19 | 1.07 | unknown protein |  |
| AT2G20570 | 1.19 | 1.39 | GBF's pro-rich region-interacting factor 1 |  |
| AT3G59190 | 1.19 | 1.46 | F-box/RNI-like superfamily protein |  |
| AT4G18140 | 1.11 | 0.89 | SCP1-like small phosphatase 4b |  |
| AT5G08430 | 1.11 | 0.92 | SWIB/MDM2 domain;Plus-3;GYF |  |
| AT3G18980 | 1.10 | 0.84 | EIN2 targeting protein1 |  |
| AT5G58340 | 1.07 | 0.87 | myb-like HTH transcriptional regulator family protein |  |
| AT1G32230 | 1.07 | 0.88 | WWE protein-protein interaction domain protein family | GO:0009414 |
|  |  |  |  | GO:0009651 |
| AT3G10450 | 1.05 | 1.18 | serine carboxypeptidase-like 7 |  |
| AT1G54370 | 1.05 | 1.04 | sodium hydrogen exchanger 5 |  |
| AT4G30900 | 1.04 | 1.05 | DNAse I-like superfamily protein |  |
| AT5G50760 | 1.00 | 0.86 | SAUR-like auxin-responsive protein family |  |
| AT1G62360 | 1.00 | 0.84 | KNOX/ELK homeobox transcription factor |  |
| AT3G24450 | 0.98 | 1.16 | Heavy metal transport/detoxification superfamily protein |  |
| AT3G51120 | 0.94 | 0.92 | DNA binding;zinc ion binding;nucleic acid binding;nucleic acid binding |  |
| AT5G67170 | 0.93 | 0.91 | SEC-C motif-containing protein / OTU-like cysteine protease family protein |  |
| AT5G38380 | 0.92 | 0.90 | unknown protein |  |
| AT4G16770 | 0.91 | 0.83 | 2-oxoglutarate (2OG) and Fe(II)-dependent oxygenase superfamily protein |  |
| AT5G42370 | 0.90 | 0.70 | Calcineurin-like metallo-phosphoesterase superfamily protein |  |
| AT1G69690 | 0.86 | 0.95 | TCP family transcription factor |  |
| AT3G53180 | 0.80 | 0.95 | glutamate-ammonia ligases;catalytics;glutamate-ammonia ligases |  |
| AT1G49360 | 0.79 | 0.75 | F-box family protein |  |
