## Supplementary material for "*DROL1* subunit of U5 snRNP in the spliceosome is specifically required to splice AT–AC-type introns in *Arabidopsis*": Table S3

Table S3. Gene Ontology analysis of upregulated genes in *drol1* mutants

| ontology | number in reference | number in list | expected | fold enrichment | p-value  |
| --- | --- | --- | --- | --- | --- |
| seed oilbody biogenesis (GO:0010344) | 9 | 4 | 0.08 | 49.82 | 1.15e-2 |
| seed development (GO:0048316) | 733 | 22 | 6.54 | 3.36 | 3.27e-3 |
| post-embryonic development (GO:0009791) | 1500 | 35 | 13.38 | 2.62 | 7.77e-4 |
| fruit development (GO:0010154) | 760 | 22 | 6.78 | 3.24 | 5.80e-3 |
| reproductive structure development (GO:0048608) | 1220 | 30 | 10.88 | 2.76 | 2.12e-3 |
| reproductive system development (GO:0061458) | 1222 | 30 | 10.90 | 2.75 | 2.20e-3 |
| response to superoxide (GO:0000303) | 13 | 4 | 0.12 | 34.49 | 3.71e-2 |
| response to oxygen radical (GO:0000305) | 13 | 4 | 0.12 | 34.49 | 3.71e-2 |
| response to stress (GO:0006950) | 3186 | 57 | 28.42 | 2.01 | 1.01e-3 |
| response to stimulus (GO:0050896) | 5464 | 92 | 48.75 | 1.89 | 5.60e-7 |
| response to oxygen-containing compound (GO:1901700) | 1431 | 45 | 12.77 | 3.52 | 8.30e-10 |
| response to chemical (GO:0042221) | 2535 | 64 | 22.62 | 2.83 | 6.31e-11 |
| response to inorganic substance (GO:0010035) | 727 | 28 | 6.49 | 4.32 | 5.34e-7 |
| lipid storage (GO:0019915) | 28 | 6 | 0.25 | 24.02 | 1.41e-3 |
| maintenance of location (GO:0051235) | 75 | 9 | 0.67 | 13.45 | 1.69e-4 |
| response to abscisic acid (GO:0009737) | 461 | 28 | 4.11 | 6.81 | 1.64e-11 |
| response to hormone (GO:0009725) | 1234 | 38 | 11.01 | 3.45 | 1.37e-7 |
| response to endogenous stimulus (GO:0009719) | 1264 | 38 | 11.28 | 3.37 | 2.69e-7 |
| response to organic substance (GO:0010033) | 1639 | 44 | 14.62 | 3.01 | 2.53e-7 |
| response to lipid (GO:0033993) | 781 | 34 | 6.97 | 4.88 | 1.83e-10 |
| response to alcohol (GO:0097305) | 466 | 28 | 4.16 | 6.74 | 2.11e-11 |
| detoxification (GO:0098754) | 153 | 9 | 1.36 | 6.59 | 4.26e-2 |
| response to toxic substance (GO:0009636) | 176 | 10 | 1.57 | 6.37 | 1.91e-2 |
| response to water deprivation (GO:0009414) | 367 | 19 | 3.27 | 5.80 | 5.67e-6 |
| response to water (GO:0009415) | 376 | 19 | 3.35 | 5.66 | 8.28e-6 |
| response to acid chemical (GO:0001101) | 408 | 19 | 3.64 | 5.22 | 2.93e-5 |
| response to abiotic stimulus (GO:0009628) | 2061 | 44 | 18.39 | 2.39 | 2.31e-4 |
| response to salt stress (GO:0009651) | 453 | 20 | 4.04 | 4.95 | 2.79e-5 |
| response to osmotic stress (GO:0006970) | 535 | 23 | 4.77 | 4.82 | 3.37e-6 |
