## Supplementary material for "*DROL1* subunit of U5 snRNP in the spliceosome is specifically required to splice AT–AC-type introns in *Arabidopsis*": Table S4

Table S5. Downregulated genes in *drol1* seedlings at 3 days after germination

| Gene ID | logFC | | Gene function | GO ID |
| --- | --- | --- | --- | --- |
|  | *drol1-1* | *drol1-2* |  |  |
| AT5G23990 | -8.70 | -5.49 | ferric reduction oxidase 5 |  |
| AT4G36570 | -7.22 | -3.10 | RAD-like 3 |  |
| AT3G24230 | -7.10 | -3.25 | Pectate lyase family protein |  |
| AT1G56100 | -6.50 | -3.80 | Plant invertase/pectin methylesterase inhibitor superfamily protein |  |
| AT5G10230 | -6.04 | -4.18 | annexin 7 |  |
| AT4G12550 | -5.93 | -3.41 | Auxin-Induced in Root cultures 1 |  |
| AT5G33355 | -5.72 | -3.86 | Defensin-like (DEFL) family protein |  |
| AT3G29410 | -5.69 | -2.47 | Terpenoid cyclases/Protein prenyltransferases superfamily protein |  |
| AT5G23980 | -5.58 | -5.41 | ferric reduction oxidase 4 |  |
| AT3G59930 | -5.43 | -3.64 | unknown protein |  |
| AT1G33670 | -5.42 | -3.27 | Leucine-rich repeat (LRR) family protein |  |
| AT5G56080 | -4.85 | -2.84 | nicotianamine synthase 2 |  |
| AT3G46900 | -4.80 | -3.89 | copper transporter 2 |  |
| AT1G71200 | -4.79 | -2.40 | basic helix-loop-helix (bHLH) DNA-binding superfamily protein |  |
| AT5G05280 | -4.74 | -1.85 | RING/U-box superfamily protein |  |
| AT5G05282 | -4.74 | -1.85 | conserved peptide upstream open reading frame 64 |  |
| AT1G52770 | -4.63 | -3.45 | Phototropic-responsive NPH3 family protein |  |
| AT3G46400 | -4.51 | -3.31 | Leucine-rich repeat protein kinase family protein |  |
| AT2G12660 | -4.49 | -3.30 | pseudogene |  |
| AT1G16910 | -4.42 | -6.49 | Protein of unknown function (DUF640) |  |
| AT3G29630 | -4.30 | -3.98 | UDP-Glycosyltransferase superfamily protein |  |
| AT4G18510 | -4.27 | -2.12 | CLAVATA3/ESR-related 2 |  |
| AT1G17300 | -4.25 | -2.13 | unknown protein |  |
| AT1G71692 | -4.24 | -2.51 | AGAMOUS-like 12 |  |
| AT1G06830 | -4.19 | -2.29 | Glutaredoxin family protein |  |
| AT3G44710 | -4.17 | -2.61 | Plant protein of unknown function (DUF247) |  |
| AT2G18800 | -4.11 | -4.04 | xyloglucan endotransglucosylase/hydrolase 21 |  |
| AT4G25010 | -4.06 | -3.05 | Nodulin MtN3 family protein |  |
| AT4G13235 | -3.99 | -1.64 | embryo sac development arrest 21 |  |
| AT2G35075 | -3.97 | -2.10 | unknown protein |  |
| AT1G22150 | -3.91 | -3.55 | sulfate transporter 1;3 |  |
| AT2G47880 | -3.85 | -3.50 | Glutaredoxin family protein |  |
| AT4G10850 | -3.80 | -4.74 | Nodulin MtN3 family protein |  |
| AT1G73410 | -3.77 | -2.71 | myb domain protein 54 |  |
| AT3G11430 | -3.74 | -2.96 | glycerol-3-phosphate acyltransferase 5 |  |
| AT4G13290 | -3.73 | -3.61 | cytochrome P450, family 71, subfamily A, polypeptide 19 |  |
| AT1G64910 | -3.69 | -3.81 | UDP-Glycosyltransferase superfamily protein |  |
| AT5G47450 | -3.53 | -2.87 | tonoplast intrinsic protein 2;3 |  |
| AT3G09220 | -3.49 | -3.39 | laccase 7 |  |
| AT2G27550 | -3.48 | -2.24 | centroradialis |  |
| AT4G15290 | -3.48 | -2.74 | Cellulose synthase family protein |  |
| AT5G09520 | -3.46 | -3.47 | hydroxyproline-rich glycoprotein family protein |  |
| AT5G50800 | -3.46 | -2.04 | Nodulin MtN3 family protein |  |
| AT4G12940 | -3.45 | -3.02 | unknown protein |  |
| AT1G63910 | -3.43 | -2.13 | myb domain protein 103 |  |
| AT1G30840 | -3.36 | -2.49 | purine permease 4 |  |
| AT2G21200 | -3.35 | -1.83 | SAUR-like auxin-responsive protein family |  |
| AT3G56240 | -3.30 | -1.73 | copper chaperone |  |
| AT2G17470 | -3.28 | -2.07 | Aluminium activated malate transporter family protein |  |
| AT4G16270 | -3.25 | -1.90 | Peroxidase superfamily protein |  |
| AT1G19900 | -3.22 | -2.84 | glyoxal oxidase-related protein |  |
| AT2G18328 | -3.21 | -2.63 | RAD-like 4 |  |
| AT4G08300 | -3.17 | -4.27 | nodulin MtN21 /EamA-like transporter family protein |  |
| AT5G48430 | -3.09 | -2.52 | Eukaryotic aspartyl protease family protein |  |
| AT3G22880 | -3.07 | -2.57 | DNA repair (Rad51) family protein |  |
| AT5G59990 | -2.93 | -2.80 | CCT motif family protein |  |
| AT1G69880 | -2.92 | -1.59 | thioredoxin H-type 8 |  |
| AT5G40690 | -2.90 | -1.87 | unknown protein |  |
| AT1G04500 | -2.82 | -2.71 | CCT motif family protein |  |
| AT1G67035 | -2.82 | -1.54 | unknown protein |  |
| AT1G78320 | -2.78 | -1.77 | glutathione S-transferase TAU 23 |  |
| AT2G21220 | -2.76 | -1.96 | SAUR-like auxin-responsive protein family |  |
| ATCG00050 | -2.75 | -2.35 | ribosomal protein S16 |  |
| AT3G50610 | -2.73 | -2.55 | unknown protein |  |
| AT1G22590 | -2.72 | -1.89 | AGAMOUS-like 87 |  |
| AT4G14020 | -2.68 | -1.41 | Rapid alkalinization factor (RALF) family protein |  |
| AT3G28570 | -2.67 | -1.67 | P-loop containing nucleoside triphosphate hydrolases superfamily protein |  |
| AT5G43290 | -2.66 | -1.25 | WRKY DNA-binding protein 49 |  |
| AT5G46140 | -2.61 | -2.07 | Protein of unknown function (DUF295) |  |
| AT5G24655 | -2.57 | -2.88 | response to low sulfur 4 |  |
| AT2G05910 | -2.49 | -2.51 | Protein of unknown function (DUF567) |  |
| AT4G24265 | -2.46 | -1.41 | unknown protein |  |
| AT1G07400 | -2.46 | -1.85 | HSP20-like chaperones superfamily protein |  |
| AT1G20030 | -2.44 | -1.39 | Pathogenesis-related thaumatin superfamily protein |  |
| AT2G41730 | -2.40 | -1.96 | unknown protein |  |
| AT2G47370 | -2.36 | -1.45 | Calcium-dependent phosphotriesterase superfamily protein |  |
| AT1G61450 | -2.36 | -1.62 | unknown protein |  |
| AT5G39090 | -2.32 | -1.58 | HXXXD-type acyl-transferase family protein |  |
| AT3G12750 | -2.31 | -1.43 | zinc transporter 1 precursor |  |
| AT1G19200 | -2.29 | -1.60 | Protein of unknown function (DUF581) |  |
| AT5G59520 | -2.27 | -1.47 | ZRT/IRT-like protein 2 |  |
| AT3G04960 | -2.26 | -1.95 | Domain of unknown function (DUF3444) |  |
| AT5G52640 | -2.24 | -1.93 | heat shock protein 90.1 |  |
| AT1G18140 | -2.23 | -1.32 | laccase 1 |  |
| AT4G26220 | -2.22 | -1.39 | S-adenosyl-L-methionine-dependent methyltransferases superfamily protein |  |
| AT5G36800 | -2.22 | -1.49 | unknown protein |  |
| AT5G36710 | -2.22 | -1.49 | unknown protein |  |
| AT4G15830 | -2.18 | -1.30 | ARM repeat superfamily protein |  |
| AT5G51600 | -2.15 | -1.24 | Microtubule associated protein (MAP65/ASE1) family protein |  |
| AT2G39080 | -2.14 | -1.12 | NAD(P)-binding Rossmann-fold superfamily protein |  |
| AT1G31335 | -2.11 | -1.51 | unknown protein |  |
| AT5G25090 | -2.11 | -1.23 | early nodulin-like protein 13 |  |
| AT5G02490 | -2.11 | -1.41 | Heat shock protein 70 (Hsp 70) family protein |  |
| AT2G26760 | -2.10 | -1.23 | Cyclin B1;4 | GO:0044772 |
|  |  |  |  | GO:0000079 |
| AT4G32830 | -2.10 | -1.26 | ataurora1 |  |
| AT4G15140 | -2.10 | -1.23 | unknown protein |  |
| AT5G24380 | -2.09 | -1.16 | YELLOW STRIPE like 2 |  |
| AT3G02120 | -2.09 | -1.33 | hydroxyproline-rich glycoprotein family protein |  |
| AT4G03100 | -2.09 | -1.22 | Rho GTPase activating protein with PAK-box/P21-Rho-binding domain |  |
| AT2G42110 | -2.08 | -1.24 | unknown protein |  |
| AT2G21640 | -2.06 | -1.63 | unknown protein |  |
| AT4G17240 | -2.05 | -1.23 | unknown protein |  |
| AT1G44110 | -2.05 | -1.16 | Cyclin A1;1 | GO:0044772 |
|  |  |  |  | GO:0000079 |
| AT2G33400 | -2.04 | -1.19 | unknown protein |  |
| AT5G16250 | -2.01 | -1.23 | unknown protein |  |
| AT4G16880 | -2.01 | -1.51 | Leucine-rich repeat (LRR) family protein |  |
| AT5G66230 | -2.01 | -1.12 | Chalcone-flavanone isomerase family protein |  |
| AT4G12900 | -2.00 | -1.62 | Gamma interferon responsive lysosomal thiol (GILT) reductase family protein |  |
| AT2G47920 | -1.99 | -1.40 | Kinase interacting (KIP1-like) family protein |  |
| AT1G47395 | -1.98 | -1.97 | unknown protein |  |
| AT1G08560 | -1.95 | -1.09 | syntaxin of plants 111 |  |
| AT5G06150 | -1.95 | -1.12 | Cyclin family protein | GO:0044772 |
|  |  |  |  | GO:0000079 |
| AT5G51440 | -1.94 | -1.49 | HSP20-like chaperones superfamily protein |  |
| AT3G18010 | -1.94 | -1.23 | WUSCHEL related homeobox 1 |  |
| AT5G45700 | -1.94 | -1.30 | Haloacid dehalogenase-like hydrolase (HAD) superfamily protein |  |
| AT1G05440 | -1.93 | -1.22 | C-8 sterol isomerases |  |
| AT1G68585 | -1.92 | -1.33 | unknown protein |  |
| AT4G31840 | -1.92 | -1.14 | early nodulin-like protein 15 |  |
| AT1G61580 | -1.92 | -1.89 | R-protein L3 B |  |
| AT3G08920 | -1.92 | -1.15 | Rhodanese/Cell cycle control phosphatase superfamily protein |  |
| AT5G36120 | -1.91 | -1.39 | cofactor assembly, complex C (B6F) |  |
| AT4G02800 | -1.91 | -1.10 | unknown protein |  |
| AT3G12870 | -1.90 | -1.15 | unknown protein |  |
| AT1G20930 | -1.89 | -1.05 | cyclin-dependent kinase B2;2 | GO:0044772 |
| AT3G52110 | -1.89 | -1.01 | unknown protein |  |
| AT2G47015 | -1.89 | -1.46 | MIR408; miRNA |  |
| AT5G58240 | -1.89 | -1.62 | FRAGILE HISTIDINE TRIAD |  |
| AT3G55660 | -1.88 | -1.21 | ROP (rho of plants) guanine nucleotide exchange factor 6 |  |
| AT3G02640 | -1.88 | -1.22 | unknown protein |  |
| AT5G56120 | -1.88 | -1.18 | unknown protein |  |
| AT1G53140 | -1.87 | -1.17 | Dynamin related protein 5A |  |
| AT3G51280 | -1.87 | -1.11 | Tetratricopeptide repeat (TPR)-like superfamily protein |  |
| AT2G25060 | -1.87 | -1.13 | early nodulin-like protein 14 |  |
| AT5G38300 | -1.84 | -1.28 | unknown protein |  |
| AT1G50490 | -1.82 | -1.20 | ubiquitin-conjugating enzyme 20 |  |
| AT1G72250 | -1.82 | -1.14 | Di-glucose binding protein with Kinesin motor domain |  |
| AT5G15510 | -1.81 | -1.07 | TPX2 (targeting protein for Xklp2) protein family |  |
| AT4G23500 | -1.81 | -1.24 | Pectin lyase-like superfamily protein |  |
| AT1G53520 | -1.80 | -1.47 | Chalcone-flavanone isomerase family protein |  |
| AT5G54560 | -1.78 | -2.27 | Protein of unknown function (DUF295) |  |
| AT4G30140 | -1.76 | -1.78 | GDSL-like Lipase/Acylhydrolase superfamily protein |  |
| AT1G75640 | -1.75 | -1.19 | Leucine-rich receptor-like protein kinase family protein |  |
| AT5G67270 | -1.74 | -0.99 | end binding protein 1C |  |
| AT4G34950 | -1.74 | -1.25 | Major facilitator superfamily protein |  |
| AT1G11220 | -1.74 | -1.31 | Protein of unknown function (DUF761) |  |
| AT3G25980 | -1.74 | -1.19 | DNA-binding HORMA family protein |  |
| AT2G45450 | -1.72 | -1.29 | protein binding |  |
| AT3G15650 | -1.72 | -1.65 | alpha/beta-Hydrolases superfamily protein |  |
| AT3G11520 | -1.72 | -1.05 | CYCLIN B1;3 | GO:0044772 |
|  |  |  |  | GO:0000079 |
| AT3G51290 | -1.72 | -1.22 | Protein of unknown function (DUF630) ;Protein of unknown function (DUF632) |  |
| AT2G46630 | -1.72 | -1.29 | unknown protein |  |
| AT3G50070 | -1.70 | -1.42 | CYCLIN D3;3 | GO:0044772 |
|  |  |  |  | GO:0000079 |
| AT3G58200 | -1.70 | -1.91 | TRAF-like family protein |  |
| AT5G22300 | -1.70 | -1.09 | nitrilase 4 |  |
| AT5G62150 | -1.70 | -1.67 | peptidoglycan-binding LysM domain-containing protein |  |
| AT5G05510 | -1.69 | -0.99 | Mad3/BUB1 homology region 1 |  |
| AT1G73630 | -1.69 | -1.60 | EF hand calcium-binding protein family |  |
| AT1G63100 | -1.68 | -0.99 | GRAS family transcription factor |  |
| AT4G24175 | -1.63 | -1.17 | unknown protein |  |
| AT5G43380 | -1.63 | -1.00 | type one serine/threonine protein phosphatase 6 |  |
| AT2G32765 | -1.62 | -1.15 | small ubiquitinrelated modifier 5 |  |
| AT4G13370 | -1.61 | -1.12 | Plant protein of unknown function (DUF936) |  |
| AT5G52860 | -1.60 | -1.36 | ABC-2 type transporter family protein |  |
| AT5G40960 | -1.55 | -1.31 | Protein of unknown function (DUF 3339) |  |
| AT3G57430 | -1.53 | -1.26 | Tetratricopeptide repeat (TPR)-like superfamily protein |  |
| AT4G11820 | -1.52 | -0.87 | hydroxymethylglutaryl-CoA synthase / HMG-CoA synthase / 3-hydroxy-3-methylglutaryl coenzyme A synthase |  |
| AT1G55140 | -1.51 | -1.02 | Ribonuclease III family protein |  |
| AT4G19840 | -1.51 | -1.11 | phloem protein 2-A1 |  |
| AT2G03505 | -1.50 | -1.29 | Carbohydrate-binding X8 domain superfamily protein |  |
| AT3G14190 | -1.49 | -1.01 | unknown protein |  |
| AT2G38620 | -1.48 | -0.96 | cyclin-dependent kinase B1;2 | GO:0044772 |
| AT5G22140 | -1.47 | -1.52 | FAD/NAD(P)-binding oxidoreductase family protein |  |
| AT1G25510 | -1.47 | -0.95 | Eukaryotic aspartyl protease family protein |  |
| AT5G11950 | -1.47 | -0.88 | Putative lysine decarboxylase family protein |  |
| AT4G34510 | -1.45 | -1.36 | 3-ketoacyl-CoA synthase 17 |  |
| AT4G37490 | -1.43 | -0.90 | CYCLIN B1;1 | GO:0044772 |
|  |  |  |  | GO:0000079 |
| AT2G29550 | -1.42 | -1.04 | tubulin beta-7 chain |  |
| AT1G54575 | -1.41 | -1.15 | unknown protein |  |
| AT1G26760 | -1.40 | -1.00 | SET domain protein 35 |  |
| AT4G34730 | -1.39 | -1.04 | ribosome-binding factor A family protein |  |
| AT1G07610 | -1.39 | -1.34 | metallothionein 1C |  |
| AT3G02240 | -1.39 | -1.50 | unknown protein |  |
| AT1G13270 | -1.38 | -0.89 | methionine aminopeptidase 1B |  |
| AT1G69700 | -1.35 | -0.90 | HVA22 homologue C |  |
| AT1G75170 | -1.33 | -0.94 | Sec14p-like phosphatidylinositol transfer family protein |  |
| AT4G29360 | -1.32 | -0.87 | O-Glycosyl hydrolases family 17 protein |  |
| AT4G33520 | -1.32 | -1.04 | P-type ATP-ase 1 |  |
| AT5G42146 | -1.32 | -1.40 | unknown protein |  |
| AT1G69230 | -1.31 | -1.46 | SPIRAL1-like2 |  |
| AT5G37010 | -1.30 | -0.89 | unknown protein |  |
| AT5G16140 | -1.29 | -0.97 | Peptidyl-tRNA hydrolase family protein |  |
| AT3G27830 | -1.28 | -1.05 | ribosomal protein L12-A |  |
| AT4G14010 | -1.28 | -1.05 | ralf-like 32 |  |
| AT1G76690 | -1.23 | -0.94 | 12-oxophytodienoate reductase 2 |  |
| AT5G64620 | -1.23 | -0.94 | cell wall / vacuolar inhibitor of fructosidase 2 |  |
| AT5G63100 | -1.22 | -0.93 | S-adenosyl-L-methionine-dependent methyltransferases superfamily protein |  |
| AT4G33470 | -1.22 | -0.99 | histone deacetylase 14 |  |
| AT3G45050 | -1.21 | -0.78 | unknown protein |  |
| AT4G25370 | -1.21 | -0.88 | Double Clp-N motif protein |  |
| AT3G27850 | -1.20 | -0.98 | ribosomal protein L12-C |  |
| AT4G35730 | -1.19 | -0.90 | Regulator of Vps4 activity in the MVB pathway protein |  |
| AT2G24440 | -1.18 | -1.07 | selenium binding |  |
| AT5G10160 | -1.18 | -0.87 | Thioesterase superfamily protein |  |
| AT3G24590 | -1.16 | -0.85 | plastidic type i signal peptidase 1 |  |
| AT5G38290 | -1.14 | -0.86 | Peptidyl-tRNA hydrolase family protein |  |
| AT3G01410 | -1.12 | -1.05 | Polynucleotidyl transferase, ribonuclease H-like superfamily protein |  |
| AT2G26840 | -1.11 | -1.02 | unknown protein |  |
| AT2G41950 | -1.11 | -0.96 | unknown protein |  |
| AT2G25880 | -1.11 | -0.87 | ataurora2 |  |
| AT4G13615 | -1.10 | -0.95 | Uncharacterised protein family SERF |  |
| AT4G03150 | -1.09 | -1.03 | unknown protein |  |
| AT4G23550 | -1.09 | -1.10 | WRKY family transcription factor |  |
| AT2G29500 | -1.08 | -0.93 | HSP20-like chaperones superfamily protein |  |
| AT3G60900 | -1.07 | -1.04 | FASCICLIN-like arabinogalactan-protein 10 |  |
| AT3G28460 | -1.05 | -0.85 | methyltransferases |  |
| AT1G01880 | -1.04 | -1.05 | 5'-3' exonuclease family protein |  |
| AT2G39290 | -1.04 | -0.82 | phosphatidylglycerolphosphate synthase 1 |  |
| AT2G05990 | -1.03 | -0.89 | NAD(P)-binding Rossmann-fold superfamily protein |  |
| AT2G21130 | -1.01 | -1.24 | Cyclophilin-like peptidyl-prolyl cis-trans isomerase family protein |  |
| AT5G55710 | -0.97 | -0.81 | unknown protein |  |
| AT3G06730 | -0.96 | -0.85 | Thioredoxin z |  |
| AT1G66670 | -0.96 | -0.85 | CLP protease proteolytic subunit 3 |  |
| AT2G04530 | -0.93 | -0.78 | Metallo-hydrolase/oxidoreductase superfamily protein |  |
| AT3G18420 | -0.92 | -0.91 | Protein prenylyltransferase superfamily protein |  |
| AT3G51140 | -0.90 | -0.81 | Protein of unknown function (DUF3353) |  |
| AT1G67660 | -0.88 | -0.79 | Restriction endonuclease, type II-like superfamily protein |  |
| AT1G21440 | -0.86 | -0.88 | Phosphoenolpyruvate carboxylase family protein |  |
| AT2G02740 | -0.80 | -0.80 | ssDNA-binding transcriptional regulator |  |
| AT1G14410 | -0.79 | -0.76 | ssDNA-binding transcriptional regulator |  |
| AT5G59500 | -0.78 | -1.03 | protein C-terminal S-isoprenylcysteine carboxyl O-methyltransferases |  |
| AT5G64150 | -0.74 | -0.83 | RNA methyltransferase family protein |  |
| AT5G20000 | -0.71 | -0.83 | AAA-type ATPase family protein |  |
