## Supplementary material for "*DROL1* subunit of U5 snRNP in the spliceosome is specifically required to splice AT–AC-type introns in *Arabidopsis*": Table S5

Table S5. Gene Ontology analysis of downregulated genes in *drol1* mutants

| ontology | number in reference | number in list | expected | fold enrichment | p-value  |
| --- | --- | --- | --- | --- | --- |
| mitotic cell cycle phase transition (GO:0044772) | 48 | 8 | 0.28 | 28.61 | 3.41e-6 |
| cell cycle phase transition (GO:0044770) | 49 | 8 | 0.29 | 28.02 | 3.95e-6 |
| cell cycle process (GO:0022402) | 415 | 16 | 2.42 | 6.62 | 1.40e-5 |
| cell cycle (GO:0007049) | 454 | 17 | 2.65 | 6.43 | 6.87e-6 |
| mitotic cell cycle process (GO:1903047) | 201 | 12 | 1.17 | 10.25 | 1.31e-5 |
| mitotic cell cycle (GO:0000278) | 245 | 13 | 1.43 | 9.11 | 1.13e-5 |
| regulation of cyclin-dependent protein serine/threonine kinase activity (GO:0000079) | 54 | 6 | 0.31 | 19.07 | 3.80e-3 |
| regulation of protein serine/threonine kinase activity (GO:0071900) | 59 | 6 | 0.34 | 17.45 | 6.12e-3 |
| regulation of protein kinase activity (GO:0045859) | 73 | 6 | 0.43 | 14.11 | 1.93e-2 |
| regulation of protein phosphorylation (GO:0001932) | 85 | 6 | 0.50 | 12.12 | 4.36e-2 |
| regulation of kinase activity (GO:0043549) | 75 | 6 | 0.44 | 13.73 | 2.23e-2 |
| regulation of cyclin-dependent protein kinase activity (GO:1904029) | 54 | 6 | 0.31 | 19.07 | 3.80e-3 |
| regulation of cell cycle (GO:0051726) | 231 | 14 | 1.35 | 10.40 | 5.37e-7 |
| regulation of mitotic cell cycle (GO:0007346) | 85 | 6 | 0.50 | 12.12 | 4.36e-2 |
