## Supplementary material for "*DROL1* subunit of U5 snRNP in the spliceosome is specifically required to splice AT–AC-type introns in *Arabidopsis*": Table S6

### Table S6

Primers for 5'RACE

|  |  |
| --- | --- |
| 5RACE\_RT | CATTTAGTATGATCC |
| 5RACE\_A1 | CACATACACTTGAACATCCTC |
| 5RACE\_A2 | CATCAACCAAAGCTACCTTAGC |
| 5RACE\_S1 | TGATATCACGTTGTTTCCTTCG |
| 5RACE\_S2 | CTTCAATGCGCATCATATGA |

Primers for gene cloning

|  |  |
| --- | --- |
| attB1DROL1p4 | GGGGCAAGTTTGTACAAAAAAGCAGGCTAAACCACTATTCTCCTTAAGC |
| attB2DROL1v3 | GGGGACCACTTTGTACAAGAAAGCTGGGTACACATCCTTGTACACG |
| attB1HD2Bp | GGGGACAAGTTTGTACAAAAAAGCAGGCTGGACCATATGGTGATCCTCAA |
| attB2HD2B | GGGGACCACTTTGTACAAGAAAGCTGGGTAAGCTCTACCCTTTCCCTTG |
| attB1HD2Cp | GGGGACAAGTTTGTACAAAAAAGCAGGCTTTGCGTTATCATAGGAAATAAACT |
| attB2HD2C | GGGGACCACTTTGTACAAGAAAGCTGGGTAAGCAGCTGCACTGTGTTT |

Primers for nucleotide substitions and insertions

|  |  |
| --- | --- |
||  |
| --- | --- |
| HD2C\_3i\_5GT | TACAAAGTTGATGCATCTGATCCGTATCCTTTACTATTTGAAC |
| HD2C\_3i\_3AG | TTCCTTGACTCTATCTCAGCGAGCCTGAGGATTTGATTG |
| HD2B\_3i\_5GT | CAAATCCCCCAACATCGAGCAGTATCCTTTTTTTGATAGATTTG |
| HD2B\_3i\_GT\_anti | TAGCCAATGAAATGAACATTTGC |
| HD2B\_3i\_3AG | TTCTTGTTGATTGAGTTTCCAGGGATGACTTCACTAGTTCG |
| HD2B\_2875A | TTAAGGAATACACAACCAATAA |
| DROL1\_2050A | CGTTTAGATAATATAACTAAAGAGAC |
| DROL1\_1AATG | AATGTCGTATAATTCTTACGCCGAGATGAGTTATC |
| DROL1\_1ATGA | ATGATCGTATAATTCTTACGCCGAGATGACTTATC |
| DROL1\_2AATG | ATGTCGTATAATTCTTACGCCGAGAATGAGTTATC |
| DROL1\_2ATGA | ATGTCGTATAATTCTTACGCCGAGATGAAGTTATC |

Primers for RT-PCR

|  |  |
| --- | --- |
| HD2B\_2721S | AAGAGTTTGAGCTTTCACACAGCG |
| HD2B\_3114A | GACAACAGCTGCTCCAGCATT |
| HD2C\_3226A | GTCATCATCTTGCTTGGCTTTG |
| HD2C\_2773S | CTGTCTACGGAGATTGTCTTG |
| HD2D\_2073A | GATCTCATTTTGAGGCAAAGCAGC |
| HD2D\_1576S | GAAATCTCTCACAGAAGTTTCC |
| NRPA2\_2624S | TACATGACACTGAAAGGACTCG |
| NRPA2\_3009A | TGTGTACGAAATCTTCGCTTGC |
| NHX5\_4217A | AGAGACAACACTCCAAGTTCTG |
| NHX5\_2171S | GATTATGATGCTCGTGCTTTCC |
